# Characterizing protein-DNA binding event subtypes in ChIP-exo data

**DOI:** 10.1101/266536

**Authors:** Naomi Yamada, William K.M. Lai, Nina Farrell, B. Franklin Pugh, Shaun Mahony

**Affiliations:** Center for Eukaryotic Gene Regulation, Department of Biochemistry & Molecular Biology, The Pennsylvania State University, University Park, PA 16802.

## Abstract

**Motivation:** Regulatory proteins associate with the genome either by directly binding cognate DNA motifs or via protein-protein interactions with other regulators. Each recruitment mechanism may be associated with distinct motifs and may also result in distinct characteristic patterns in high-resolution protein-DNA binding assays. For example, the ChIP-exo protocol precisely characterizes protein-DNA crosslinking patterns by combining chromatin immunoprecipitation (ChIP) with 5’ → 3’ exonuclease digestion. Since different regulatory complexes will result in different protein-DNA crosslinking signatures, analysis of ChIP-exo tag enrichment patterns should enable detection of multiple protein-DNA binding modes for a given regulatory protein. However, current ChIP-exo analysis methods either treat all binding events as being of a uniform type or rely on motifs to cluster binding events into subtypes.

**Results:** To systematically detect multiple protein-DNA interaction modes in a single ChIP-exo experiment, we introduce the ChIP-exo mixture model (ChExMix). ChExMix probabilistically models the genomic locations and subtype memberships of binding events using both ChIP-exo tag distribution patterns and DNA motifs. We demonstrate that ChExMix achieves accurate detection and classification of binding event subtypes using *in silico* mixed ChIP-exo data. We further demonstrate the unique analysis abilities of ChExMix using a collection of ChIP-exo experiments that profile the binding of key transcription factors in MCF-7 cells. In these data, ChExMix identifies possible recruitment mechanisms of FoxA1 and ERα, thus demonstrating that ChExMix can effectively stratify ChIP-exo binding events into biologically meaningful subtypes.

**Availability:** ChExMix is available from https://github.com/seqcode/chexmix

## Introduction

Sequence-specific transcription factors (TFs) recognize many of their regulatory targets by making direct contact with their cognate DNA binding sites. However, TFs and other regulatory proteins can also associate with DNA indirectly, via protein-protein interactions with cooperating DNA-bound regulators. Genome-wide protein-DNA interaction assays such as ChIP-seq (Barski et al., 2007; Johnson et al., 2007) and ChIP-exo (Rhee and Pugh, 2011) typically rely on agents that induce both protein-DNA and protein-protein crosslinking, and therefore do not necessarily discriminate between such direct and indirect DNA binding modes. In fact, some studies report that up to two thirds of *in vivo* TF binding events, defined here as precise locations where the TF associates with the genome, lack cognate motif instances (Wang et al., 2012; Starick et al., 2015). Hence, a single ChIP-seq or ChIP-exo experiment might encompass diverse binding event types, produced by different protein-DNA interaction modes.

ChIP-exo and related assays (e.g. ChIP-nexus (He et al., 2015)) precisely define protein-DNA crosslinking patterns with the use of lambda exonuclease (Rhee and Pugh, 2011). The exonuclease digests DNA in a 5’ to 3’ direction and, on average, stops at 6bp before a protein-DNA crosslinking point. Since different regulatory complexes will result in different crosslinking signatures, analysis of ChIP-exo sequencing tag distribution patterns around a given protein’s DNA binding events should enable detection of multiple protein-DNA binding modes. For example, Starick, *et al.* characterized glucocorticoid receptor (GR) binding using ChIP-exo and classified detected binding events using motif information. This approach uncovered a subset of GR ChIP-exo peaks that contained a Forkhead TF DNA binding motif (Starick et al., 2015). The same sites displayed a distinct ChIP-exo tag distribution pattern from that observed at peaks containing the GR cognate binding motif. The authors thereby hypothesized that some ChIP-exo derived GR binding events represent indirect binding to DNA via protein-protein interactions with a Forkhead TF. Therefore, careful analysis of ChIP-exo tag distribution patterns and DNA binding motifs may enable discrimination between a protein’s distinct DNA binding modes.

Most available approaches for discriminating between direct and indirect binding modes in a ChIP-seq or ChIP-exo experiment rely exclusively on DNA motif analysis. For example, several methods assume that directly bound sites should contain an instance of a cognate binding motif, while indirectly bound sites will contain motif instances corresponding to other TFs (Bailey and MacHanick, 2012; Whitington et al., 2011; Gordân et al., 2009; Neph et al., 2012; Keilwagen and Grau, 2015). This assumption may not always be true. Distinct regulatory complexes may not always be associated with distinct DNA binding motifs, although they may still be distinguishable based on variations in ChIP crosslinking patterns. Therefore, analyzing combinations of both DNA sequence and ChIP tag distribution information may be necessary to fully characterize the diversity of protein-DNA binding modes present in a given experiment.

One previous approach has attempted to cluster TF binding events using ChIP-seq tag enrichment patterns, and reports on each cluster’s associations with GO terms, motif enrichment, genomic localization, and gene expression (Cremona et al., 2015). However, clustering ChIP-seq tag enrichment patterns is confounded by high variance in the locations of ChIP-seq tags with respect to the protein-DNA binding event. ChIP-seq resolution is limited by sonication, which results in broad tag distributions. As described above, the ChIP-exo assay is more appropriate for characterizing distinct binding modes via analysis of tag distribution shapes, because ChIP-exo tag distributions are determined by crosslinking patterns at each binding site. However, no available method can exploit tag distribution patterns to delineate distinct protein-DNA binding modes in a ChIP-exo experiment.

To systematically detect multiple protein-DNA interaction modes in a single ChIP-exo experiment, we introduce the ChIP-exo mixture model (ChExMix). ChExMix discovers and characterizes binding event subtypes in ChIP-exo data by leveraging both sequencing tag enrichment patterns and DNA motifs. In doing so, ChExMix offers a more principled and robust approach to characterizing binding subtypes than simply clustering binding events using motif information. For instance, ChExMix does not require that all (or any) subtype-specific binding events be associated with motif instances, thus enabling binding subtype classification only using ChIP-exo tag patterns.

To demonstrate its unique analysis abilities, we applied ChExMix to ChIP-exo data profiling key regulators in estrogen receptor (ER) positive breast cancer cells. Upon estradiol treatment, FoxA1, ERα, and CTCF co-localize at a subset of genomic locations. Our findings suggest that FoxA1 likely binds to some genomic loci via protein-protein interactions with ERα and CTCF. Conversely, indirect binding of ERα to DNA via FoxA1 interactions is also observed in ERα ChIP-exo. These results demonstrate that ChExMix can characterize multiple protein-DNA interaction modes in ChIP-exo data, providing us with unique insights into interactions between transcription factors in a given cell type.

## Results

### ChExMix model overview

ChExMix integrates information from ChIP-exo tag distributions and DNA sequences in a probabilistic mixture model framework to characterize multiple DNA-protein interaction modes. Initial candidate ChIP-exo peak locations are determined using a probabilistic mixture model that doesn’t incorporate subtypes, similar to the approach described in our previously published GPS ChIP-seq peak-finder (Guo et al., 2010) (Figure 1A). Using these initial binding event locations, ChExMix estimates potential subtypes by performing *de novo* motif discovery around the predicted binding events and/or by clustering tag distributions in 150bp windows using Affinity Propagation (Figure 1B). Discovered subtypes that have similar motifs and tag distributions are merged. Lastly, ChExMix assigns binding events to subtypes using a hierarchical mixture model (Figure 1C). ChExMix probabilistically assigns observed tags to binding events by calculating the probabilities that each tag was generated by each binding event given the binding events’ current locations, strengths (mixing probabilities), subtype assignments, and the tag distributions associated with each subtype. The Expectation Maximization (EM) algorithm is used to iteratively optimize the positions, strengths, and subtype membership of each binding event using information from both the assigned tags and the underlying DNA sequences. In estimating the subtype probabilities for each binding event, we incorporate the following biologically-motivated assumptions in the form of priors: 1) a sparseness prior biases the algorithm to associate each binding event with a single binding subtype; and 2) the presence of a particular subtype’s motif at a binding event biases the assignment of the binding event to that subtype. ChExMix takes mapped tags (e.g. BAM files) as input and outputs binding event positions and subtype assignments. ChExMix runs within a few hours for most datasets (Table S1).

**Figure 1.**
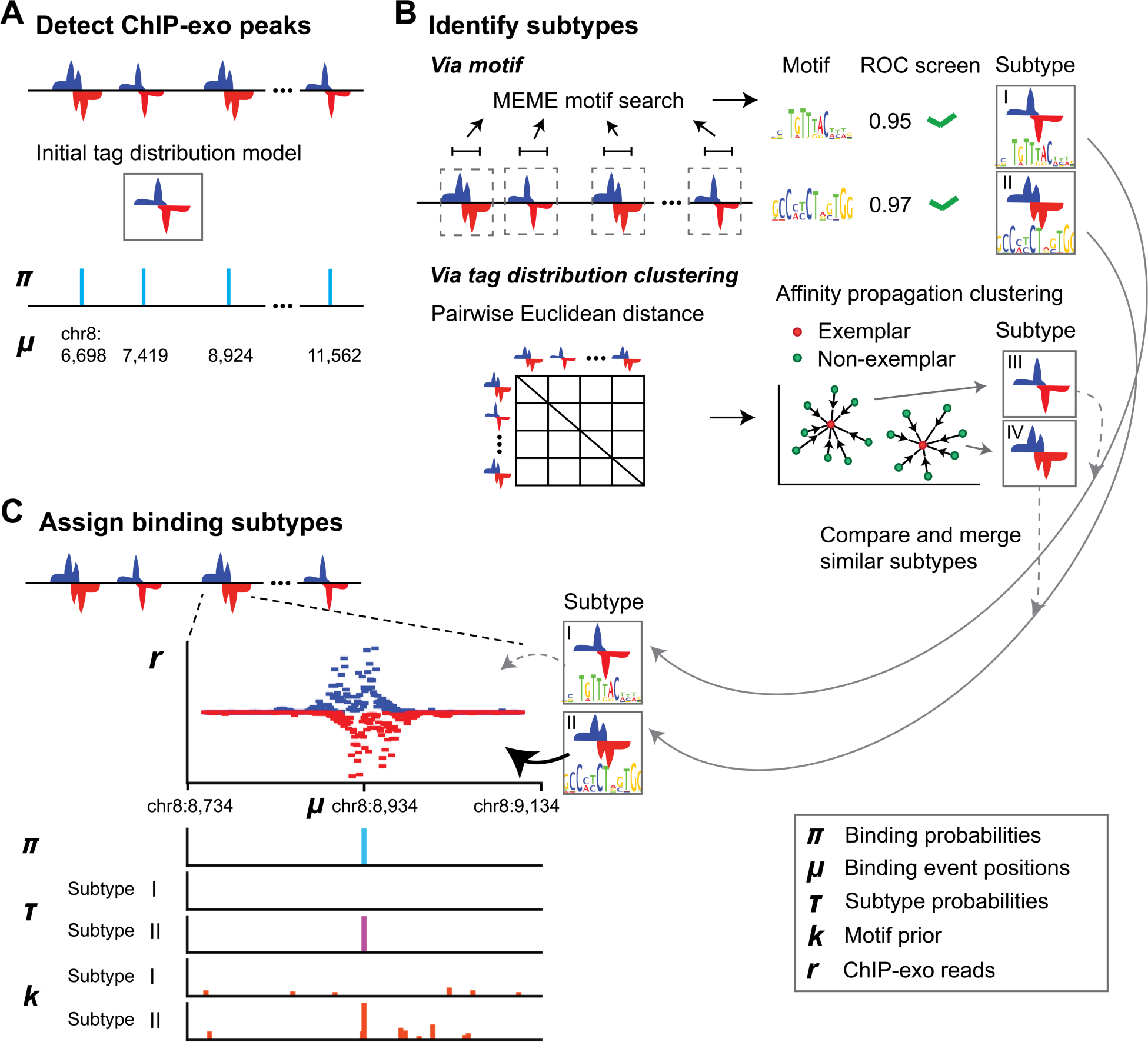
Overview of the ChExMix model. A) ChExMix first detects ChIP-exo peaks genome-wide using a probabilistic mixture model that does not incorporate subtypes. B) ChExMix defines subtypes via motif discovery and/or by clustering tag patterns around predicted binding events. C) ChExMix uses a hierarchical mixture model to assign binding events to subtypes and to optimize their locations. The illustration shows an example of final ChExMix parameter values at a binding event location.

### ChExMix accurately classifies binding subtypes in *in silico* mixed ChIP-exo datasets

ChExMix is designed to discover and model multiple binding subtypes within a single ChIP-exo dataset. We cannot assume *a priori* that we know the correct assignment of TF binding events to subtypes in any existing ChIP-exo experiment. Therefore, to test the ability of ChExMix to estimate binding subtypes and assign binding events to subtypes, we created datasets that mix data from two distinct ChIP-exo experiments (and thus contain definitive assignments of binding events to two distinct “subtypes”).

Specifically, we computationally mixed ChIP-exo data from CTCF and FoxA1, two TFs that are known to produce distinct ChIP-exo tag distribution patterns at their respective binding events (Rhee and Pugh, 2011; Serandour et al., 2013). The locations of binding events in the mixed experiments were defined by selecting equal numbers of non-overlapping binding events for each TF (see Methods). The signal portion of our mixed experiments was then defined by randomly selecting CTCF ChIP-exo tags from the CTCF binding event locations and FoxA1 ChIP-exo tags from the FoxA1 binding event locations. Each simulated experiment contains 6 million signal tags, but the relative frequency at which CTCF and FoxA1 tags were selected was varied to simulate subtypes having different relative representations in a dataset. A further set of 24 million background tags were drawn at random from a control (input) experiment.

In the simulated setting in which there is equal representation of CTCF and FoxA1 subtypes (i.e. 3 million tags drawn from each dataset), ChExMix discovers two distinct subtypes characterized by subtype-specific DNA motifs and tag distributions associated with CTCF (Figure 2A) and FoxA1 (Figure 2B). ChExMix also achieves high performance in appropriately assigning binding events to their source CTCF and FoxA1 “subtypes” (CTCF: Figure 2C red dots, TPR=88.9%, FPR=3.5%; FoxA1: Figure 2D red dots, TPR=96.5%, FPR=11.1%; Figure S1A,B; Figure S2A,B; Table S2). ChExMix performance in detecting the two subtypes and appropriately assigning subtypes to binding events remains high over all relative sampling rates tested from the CTCF and FoxA1 subtypes, suggesting that subtypes do not have to be present in equal proportions in order for ChExMix to discover them. ChExMix also maintains high performance over various read depths (Figure S3), biological replicates (Figure S4), and simulation setting where subtypes either have different motifs (Figure S5; Figure S6) or tag distributions (Figure S7), but not both.

**Figure 2.**
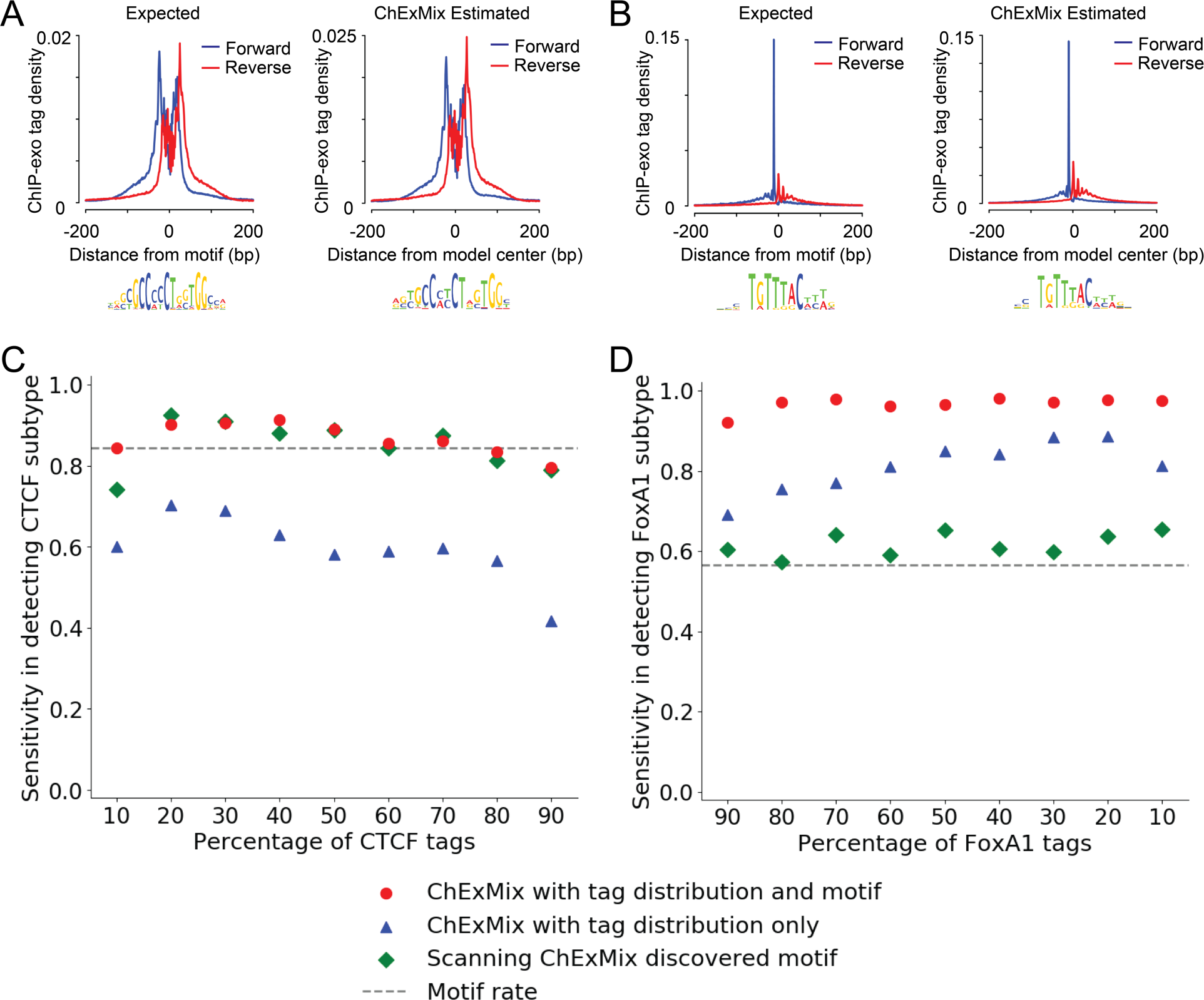
ChExMix learns subtype-specific tag distributions and accurately predicts binding event subtypes in *in silico* mixed CTCF and FoxA1 ChIP-exo data. **A)** CTCF ChIP-exo tag distribution (forward strand in blue and reverse strand in red) at CTCF motif locations (left). CTCF subtype-specific tag distribution model and motif learned by ChExMix (right). **B)** FoxA1 ChIP-exo tag distribution (forward strand in blue and reverse strand in red) at FoxA1 motif locations (left). FoxA1 subtype-specific tag distribution model and motif learned by ChExMix (right). **C), D)** Sensitivity in subtype assignment using ChExMix with *de novo* estimated tag distributions and motifs (red dots) and ChExMix with tag distributions alone (blue triangles). Fraction of peaks containing ChExMix discovered motifs (green diamonds). Plots show sensitivity for correctly assigning binding events to the CTCF (C) and FoxA1 (D) subtypes, as the relative proportion of signal tags is varied between the CTCF and FoxA1 experiments. Each data point represents an average performance over five simulated datasets (see Figure S1). Matching specificity plots in Figure S2.

By uniquely combining both DNA motifs and ChIP-exo tag distributions to classify binding subtypes, ChExMix outperforms alternative approaches that use one or the other source of information in subtype assignment. For example, a motif scanning approach that classifies binding events based on the presence of ChExMix discovered motifs fails to appropriately classify many of the FoxA1 subtype binding events (Figure 2D green diamonds; Figure S1E,F). Similarly, a version of ChExMix that uses only tag information in subtype assignment (subtypes are still defined using both motif discovery and tag distributions) displays lower sensitivity than the version of ChExMix that uses both tag distributions and DNA motifs (Figure 2C blue triangles; Figure S1C,D; Table S2). Our results thus demonstrate that ChExMix enables discovery of binding subtypes within a single ChIP-exo dataset and accurately assigns subtypes to binding events.

### ChExMix enables discovery of binding subtypes using only ChIP-exo tag distributions

ChExMix’s combined use of DNA motifs and ChIP-exo tag distributions has obvious advantages when the regulatory protein of interest is a sequence-specific TF. However, characterizing and classifying binding event subtypes may also be useful in the analysis of regulatory proteins that lack an obvious sequence preference. ChExMix can characterize binding subtypes without any sequence motif information by clustering binding event ChIP-exo tag distributions using Affinity Propagation (Dueck and Frey, 2007). To demonstrate that ChExMix can thereby discover and assign *de novo* binding subtypes using only tag distribution information, we assessed its performance in a controlled simulation setting where no specific sequence signals were introduced.

We simulated 500 binding events from each of two distinct types by randomly drawing tags from two pre-defined ChIP-exo distribution patterns (Figure 3A, 3B; see Methods). The 1,000 binding events were placed at defined locations along the yeast genome. Each simulated experiment contains 100, 200, and 300 thousand signal tags (i.e. drawn from the ChIP-exo distributions in proximity to one of the simulated binding events). The relative frequency at which each of the two subtypes’ tags were selected was varied to simulate subtypes having different representations in a dataset. Further sets of background tags were drawn from a yeast control (mock IP) experiment, resulting in a total of one million tags per simulation dataset.

In the simulated setting in which there is equal representation of both subtypes (and 20% of tags are sampled from signal regions), ChExMix successfully recovers the two distinct subtypes by clustering the initial binding events (Figure 3C, 3D). During ChExMix training, the two estimated subtype tag distributions are further refined (Figure 3E, 3F), and the end results closely resemble the original distributions (Figure 3A, 3B). ChExMix achieves high performance in appropriately assigning binding events to the two subtypes (Subtype A: Figure 3G orange dots, TPR=99.8%, FPR=5.9%; Subtype B: Figure 3H orange dots, TPR=94.1%, FPR=0.2%). ChExMix maintains this high performance in detecting and assigning subtypes in cases where one of the subtypes has a relatively low representation in the dataset, or when the overall signal in the ChIP-exo experiment is relatively low (Figure 3G, 3H; Figure S8). The simulation experiments thus demonstrate that ChExMix has the unique ability to accurately identify and assign binding event subtypes even if no distinctive DNA motifs are associated with those subtypes.

**Figure 3.**
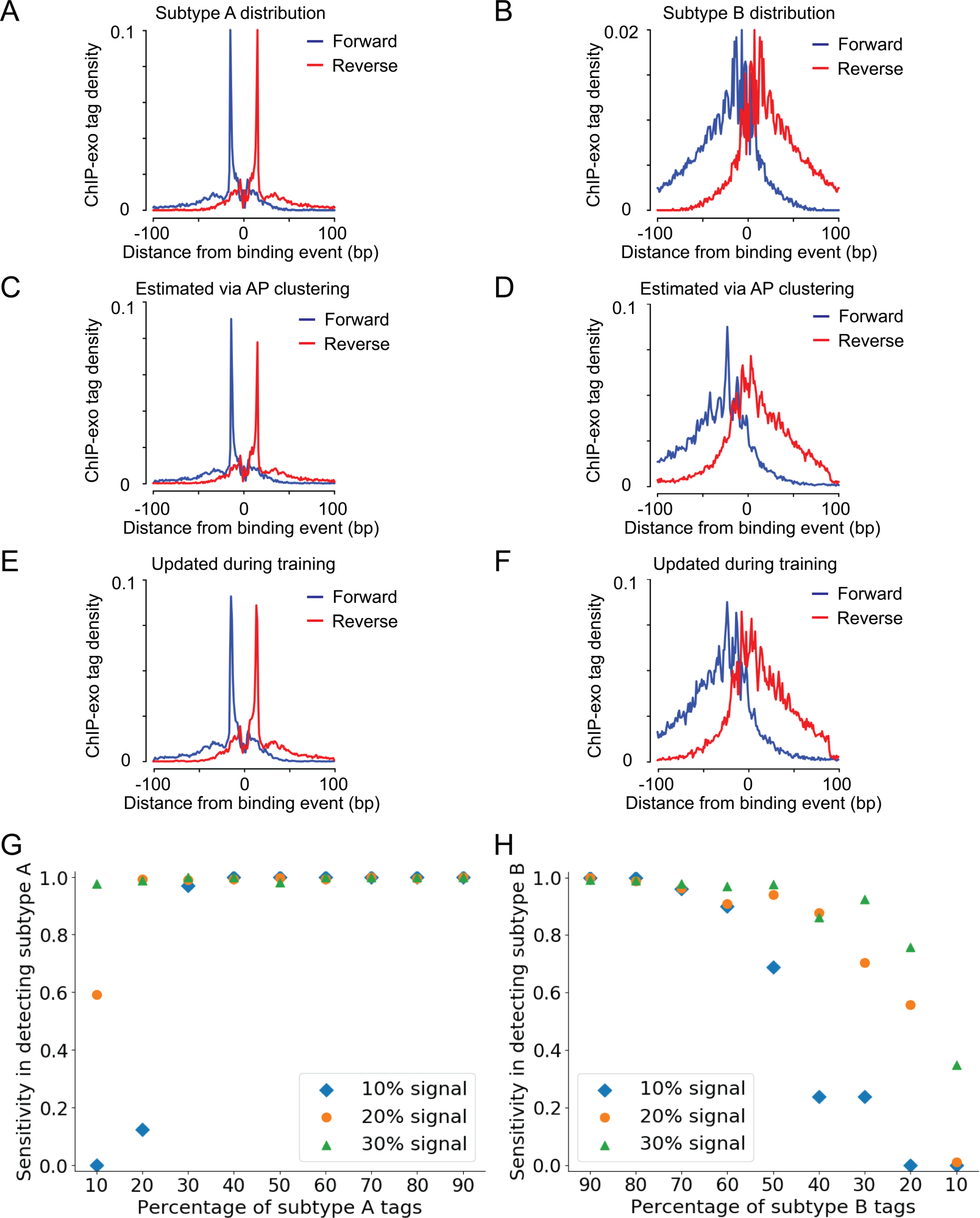
ChExMix learns subtype specific tag distributions *de novo* and accurately predicts binding event subtypes without motif information. **A), B)** Simulation data contains binding events from two distinct subtypes that have distinct tag distributions. **C), D)** In the 20% signal simulation setting, ChExMix appropriately discovers two distinct distributions via affinity propagation clustering. **E), F)** The *de novo* discovered distributions are further refined during ChExMix training. The 5’ ends of forward and reverse strand tags are shown in blue and red lines, respectively. **G), H)** Sensitivity in subtype assignment using *de novo* estimated tag distributions with overall signal of 10% (blue diamonds), 20% (orange dots), and 30% (green triangles). Plots show sensitivity for correctly assigning binding events to the subtype A (Reb1 distribution) (G) and subtype B (p53 distribution) (H) subtypes, as the relative proportion of signal tags is varied between the two subtypes.

### ChExMix maintains high accuracy in predicting binding event locations

We have previously demonstrated that the probabilistic mixture modeling framework underlying GPS, GEM, and MultiGPS enables highly accurate protein-DNA binding event detection in ChIP-seq and ChIP-exo data (Guo et al., 2010, 2012; Mahony et al., 2014). Since ChExMix substantially modifies this framework to account for binding event subtypes, we assessed whether these changes have negatively impacted the ability to characterize binding locations.

We compared ChExMix performance in predicting human CTCF (Rhee and Pugh, 2011) and mouse FoxA2 (Iwafuchi-Doi et al., 2016) binding event locations to that of nine ChIP-exo analysis methods, including MultiGPS (Mahony et al., 2014), GEM (Guo et al., 2012), MACS2 (Zhang et al., 2008), MACE (Wang et al., 2014), PeakXus (Hartonen et al., 2016), Peakzilla (Bardet et al., 2013), Q-nexus (Hansen et al., 2016), DFilter (Kumar et al., 2013), and CexoR (Madrigal, 2015). We excluded ChIP-ePENS (Ye et al., 2016) from our evaluation because it requires paired-end ChIP-exo data. Both CTCF and FoxA2 ChIP-exo datasets consist of single-end sequencing data.

To ensure a fair comparison, we used 1,553 shared CTCF sites that are predicted by all ten methods and which contain an instance of the CTCF motif within 50bp. Spatial resolution is measured by the difference between the computationally predicted locations of binding events and the nearest match to the proximal consensus motif. Thus, by design of the comparison, all methods locate 100% of these events within 50bp of the motif position. ChExMix exactly locates the events at the motif position in 87.5% of these events, outperforming all other methods (Figure 4A). Similarly, we identified 835 FoxA2 sites in the FoxA2 ChIP-exo dataset that are predicted by nine methods excluding CexoR and which contain an instance of the FoxA2 motif within 50bp. CexoR requires replicated experiments; the FoxA2 ChIP-exo replicate has a low sequencing depth and is not adequate for CexoR analysis. ChExMix exactly located the events at the motif position in 64.0% of these events (Figure 4C). ChExMix binding event predictions also contain instances of the cognate motifs at a high rate (Figure 4B, D). Similarly, ChExMix retains high resolving power in detecting two closely placed binding events (Figure S9) as previously demonstrated in the GPS framework. These results suggest that ChExMix maintains high accuracy in protein-DNA binding event prediction.

**Figure 4.**
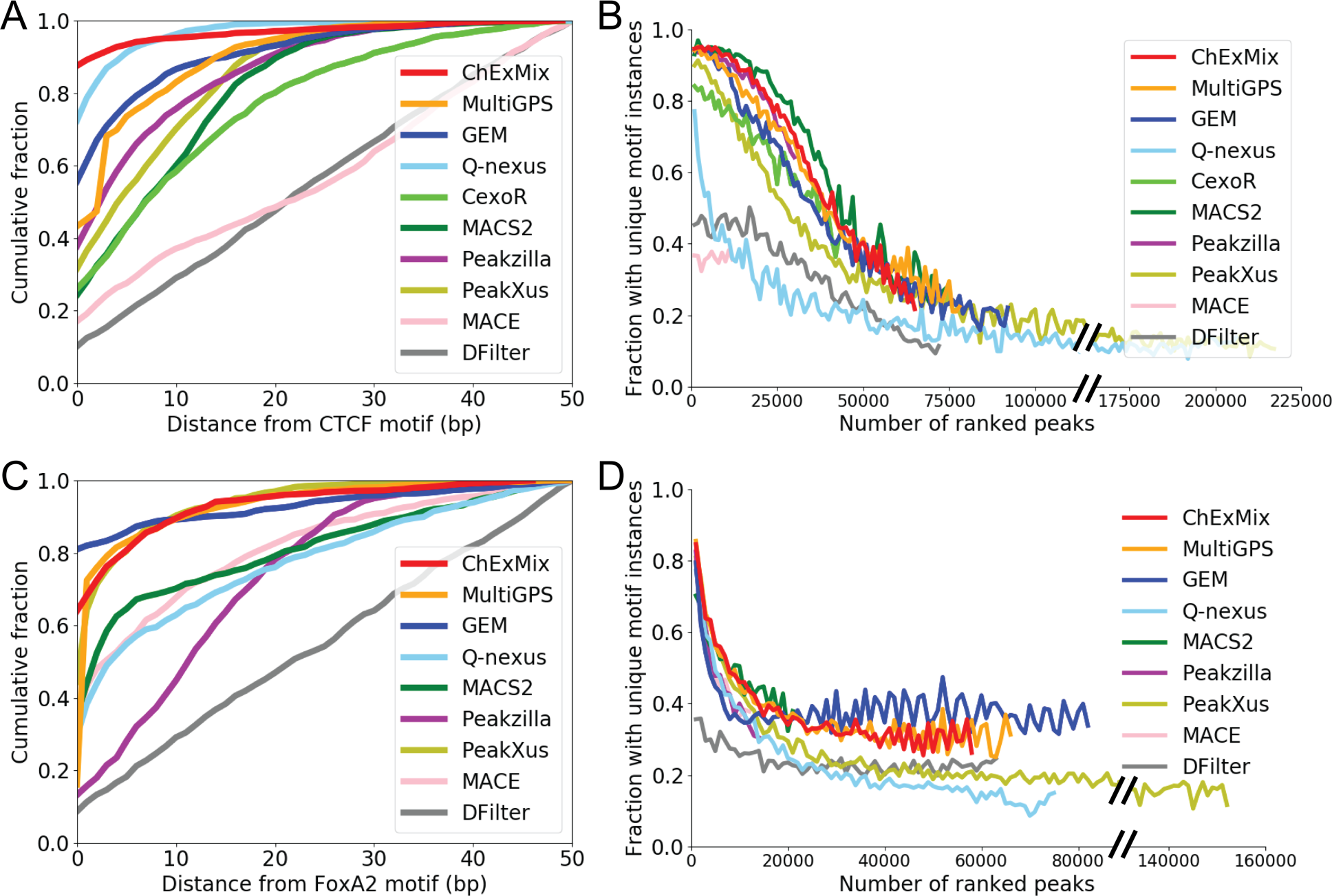
ChExMix accurately estimates binding event locations. **A)** Cumulative fraction of selected CTCF binding event predictions that have a CTCF motif instance present within the given distance following event discovery by ChExMix, MultiGPS, GEM, Q-nexus, CexoR, MACS2, Peakzilla, PeakXus, MACE, and DFilter. Events evaluated were predicted by all ten methods and had a CTCF motif instance within 50bp. **B)** Fraction of each method’s ranked CTCF binding event predictions that have a unique CTCF motif instance present within 50bp. **C)** Cumulative fraction of selected FoxA2 binding event predictions that have a FoxA2 motif instance present within the given distance following event discovery by ChExMix, MultiGPS, GEM, Q-nexus, MACS2, Peakzilla, PeakXus, MACE, and DFilter. Events evaluated were predicted by all nine methods and had a FoxA2 motif instance within 50bp. **D)** Fraction of each method’s ranked FoxA2 binding event predictions that have a unique FoxA2 motif instance present within 50bp.

### ChExMix deconvolves regulatory molecule interactions of FoxA1, Estrogen Receptor α, and CTCF in MCF-7 cells

To demonstrate the ability of ChExMix to discover biologically relevant binding event subtypes, we applied ChExMix to analyze FoxA1 ChIP-exo data in MCF-7 cells. The pioneer factor FoxA1 is a key determinant of estrogen receptor function and endocrine response, and influences genome-wide accessibility in MCF-7, thus affecting global ER binding (Hurtado et al., 2011). CTCF is an upstream negative regulator of FoxA1 and ER chromatin interactions (Hurtado et al., 2011; Fiorito et al., 2016). Genome-wide profiling suggest that these factors co-localize at a subset of binding loci, but how these factors interact with one another and DNA at specific sites remains largely unevaluated.

ChExMix identifies three main subclasses in FoxA1 ChIP-exo data. The majority (24,749) of binding events are associated with a subtype that contains FoxA1’s cognate DNA binding motif and a ChIP-exo tag distribution shape highly similar to that found in previous ChIP-exo analyses of FoxA transcription factors (Iwafuchi-Doi et al., 2016; Ye et al., 2016; Serandour et al., 2013) (Figure 5A, 5B; Figure S10A,B; Table S3). We thus label this the “direct binding” subtype. However, 2,666 binding events are assigned to subtype 1, which contains a nuclear hormone receptor DNA binding motif similar to that bound by ERα (Figure 5A). Similarly, 2,648 events are assigned to subtype 2, which contains a CTCF-like motif. Both subclasses are also associated with distinct tag distributions (Figure 5B).

**Figure 5.**
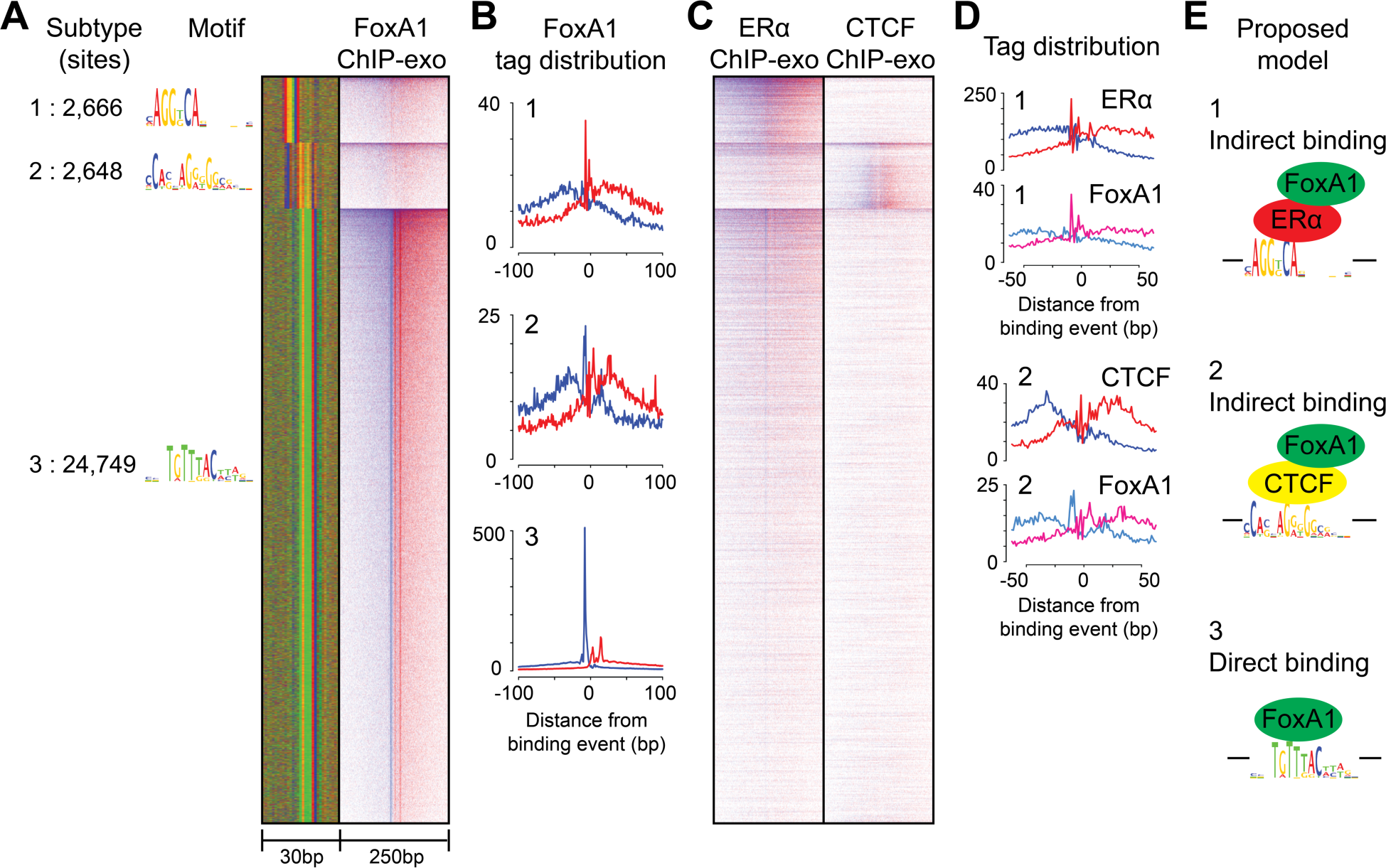
ChExMix discovers site-specific recruitment of FoxA1 via ERα and CTCF in MCF-7 FoxA1 ChIP-exo data. **A)** Motif, sequence color plot, and heatmap of three subtypes identified in FoxA1 ChIP-exo. Sites within each subtype are aligned by the ChExMix-defined binding event position and orientation. Different subtypes are aligned to each other via motif alignment. **B)** FoxA1 tag pattern associated with subclass 1, 2, and 3. **C)** Heatmaps of ERα and CTCF ChIP-exo tags at FoxA1 binding events. **D)** ERα tag pattern at subclass 1 binding events and CTCF tag pattern at subclass 2 binding events. **E)** Proposed TF interactions between FoxA1, ERα, and CTCF.

We hypothesized that subtypes 1 & 2 represent indirect FoxA1 binding to DNA via protein-protein interactions with ERα and CTCF, respectively (Figure 5E). We thus examined whether subtypes 1 & 2 are bound by their respective predicted factors using ERα and CTCF ChIP-exo datasets. We found that 55.4% of subclass 1 events are located within 100bp of ERα binding events, while 37.5% of the subclass 2 events occur within 100bp of CTCF ChIP-exo peaks (Figure 5C) (Poisson *p*-value < 0.001 for the overlap between subtype 1 and ERα binding and between subtype 2 and CTCF binding). The tag distribution shape of subtype 1 binding events in FoxA1 ChIP-exo resembles the tag distribution shape in ERα at the same sites, peaking at the exact same base positions (Figure 5D).

We further hypothesized that if FoxA1 binding is mediated via ERα at subtype 1 locations in MCF-7 cells, we should observe FoxA1 binding to fewer subtype 1 locations in ER negative breast cancer cells. In accordance with this hypothesis, only 30.4% (811/2,666) of FoxA1 subtype 1 binding events occur within 50bp of a FoxA1 ChIP-exo peak in MDA-MB-453 (an ER negative breast cancer cell line). In contrast, 59.6% (14,761/24,749) of FoxA1 subtype 3 events are bound in MDA-MB-453. These results are consistent with our hypothesis of indirect FoxA1 binding at subtype 1. We found no evidence that the various detected subtypes correspond to differences in transcriptional behavior within MCF-7 cells (Figure S11; Figure S12). The fact that the overlap of these subtypes with ERα and CTCF binding events is incomplete may be due to thresholding effects, erroneous assignments of FoxA1 binding events to the relevant subtypes, or may possibly reflect FoxA1 interactions with other transcription factors that have similar binding preferences. For example, several nuclear hormone receptors are active in MCF-7 cells, including Progesterone Receptor and Glucocorticoid Receptor, and are expected to bind to DNA binding motifs related to that discovered at subtype 1 binding events.

We next applied ChExMix to analyze ERα ChIP-exo data, discovering seven distinct subtypes (Figure 6A; Figure S10C,D; Table S3). The majority (24,914) of binding events are associated with one of six subtypes that contains a nuclear hormone receptor motif, which ERα may be expected to directly bind. However, 3,009 binding events are associated with subtype 4, which contains a Forkhead motif similar to that bound by FoxA1. Subtype 4 is also associated with a distinct tag distribution shape (Figure 6B), again suggesting a hypothesis whereby ERα binds indirectly via protein-protein interactions with FoxA1 at subtype 4 binding events (Figure 6E). Indeed, 62.8% of subclass 4 events are located within 100bp of FoxA1 binding events (Figure 6C), and the ERα ChIP-exo tag distribution at subtype 4 binding events peaks at the same base pair positions as the FoxA1 ChIP-exo tag distribution at the same sites (Figure 6D). These results strongly suggest that ChExMix can discover binding event subtypes representing direct and indirect TF interactions from a single ChIP-exo experiment.

**Figure 6.**
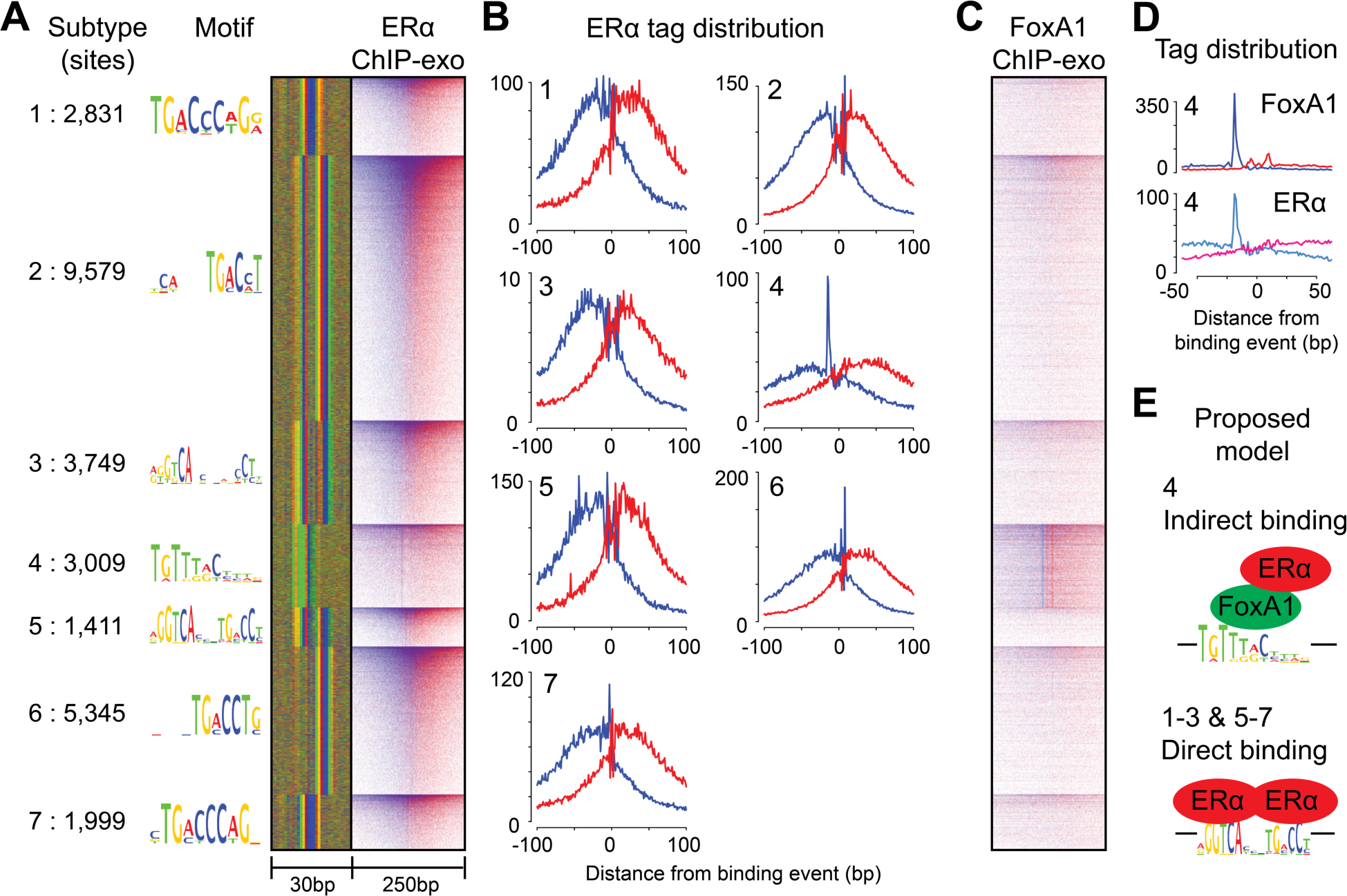
ChExMix discovers FoxAl mediated ERα binding in MCF-7 ERα ChlP-exo data. **A)** Motif, sequence plot and heatmap of seven subclasses identified in ERα ChlP-exo. Sites within each subtype are aligned by the ChExMix-defined binding event position and orientation. Different subtypes are aligned to each other via motif alignment. **B)** ERα tag patterns centered at subclass binding events. **C)** Heatmap of FoxA1 ChIP-exo tags centered at ERα predicted binding events. **D)** FoxA1 tag distribution centered at ERα subtype 4 binding events. **E)** Proposed binding models of ERα subtypes.

## Discussion

ChExMix provides a principled platform for elucidating diverse protein-DNA interaction modes in a single ChIP-exo experiment by exploiting both ChIP-exo tag enrichment patterns and DNA motifs. Using a fully integrated framework, ChExMix allows simultaneous detection of binding event locations, discovery of binding event subtypes, and assignment of binding events to subtypes. As demonstrated above, ChExMix provides highly accurate spatial resolution of binding event predictions and accurately assigns binding events to subtypes. Uniquely, ChExMix can characterize binding event subtypes without requiring the presence of distinctive sequence features, thus potentially enabling binding subtype analysis of non-sequence-specific regulatory proteins (e.g. chromatin modifiers, co-activators, co-repressors, etc.).

We further demonstrated that ChExMix can characterize biologically relevant binding event subtypes in ER positive breast cancer cells. FoxA1, ERα, and CTCF have previously been shown to co-localize at some sites, but their modes of interaction with one another remained elusive. In FoxA1 ChIP-exo data, ChExMix identifies subtypes corresponding to ERα and CTCF motifs, and about a half of these subtypes’ binding events are bound by the ERα and CTCF proteins, respectively. Our results thus suggest that ERα and CTCF likely mediate binding of FoxA1 via protein-protein interactions at a subset of the genomic loci where multiple factors are co-bound. The analysis presented in the paper is restricted to the most over-represented subtypes associated with the FoxA1 and ERα ChIP-exo datasets. Because FoxA1 and ERα have been shown to co-localize with several other transcription factors, the results presented here may not include a comprehensive set of factors with which FoxA1 and ERα interact. Future improvements of the method may include richer sequence analysis to recover motifs with lower representation, and the application of metrics to test subtype-specific motifs based on how centrally tags are enriched around the motifs. Another possible approach for discovering weaker subtypes is to initialize a large number of potential subtypes using compendia of known TF binding motifs and to rely on EM training to weed out non-significant ones.

In summary, we have demonstrated that ChExMix enables new forms of insight from a single ChIP-exo experiment, taking analysis beyond merely cataloging binding event locations and towards a fine-grained characterization of distinct protein-DNA binding modes. As demonstrated in our MCF-7 analyses, integrating ChExMix analyses across collections of related ChIP-exo experiments will enable us to identify the individual transcription factors responsible for recruiting several regulatory proteins, and thus modulating regulatory activities, at specific genomic loci.

## Methods

### ChExMix hierarchical mixture model

Similar to the previously described GPS (Guo et al., 2010), GEM (Guo et al., 2012), and MultiGPS (Mahony et al., 2014) approaches to ChIP-seq binding event detection, ChExMix models ChIP-exo sequencing data as being generated by a mixture of binding events along the genome, and an Expectation Maximization (EM) learning scheme is used to probabilistically assign sequencing tags to binding event locations. The GPS, GEM, and MultiGPS frameworks assume that a single experiment-specific tag distribution generates all binding events in a given dataset. ChExMix breaks this assumption by modeling multiple distributions within a single dataset. ChExMix further models binding events as a mixture of binding subtypes, where each subtype *t* is defined by a distinct tag distribution and possibly a distinct DNA motif. Since the tag distributions and motifs are strand-asymmetric, each subtype has an implicit orientation. To account for the expected equal representation of each binding event subtype on both DNA strands, we define the subtypes in pairs, where the tag distributions and motifs in each pair are constrained to be reverse-complements of each other.

The empirically estimated multinomial distribution Pr(*r_n_*|*x*, *t*) gives the strand-specific probability of observing ChIP-exo tag *r_n_* from a binding event of subtype *t* located at genomic coordinate *x*. We define a vector of component locations *μ* where *μ_j,t_* is the genomic location of event *j* of the binding subtype *t*. In other words, the binding event’s exact location within a genomic locus is dependent on the estimated subtype. Similarly, we introduce a vector of component subtype probabilities *τ*, where *τ_j,t_* is the probability of the binding event *j* belonging to subtype *t*. We initialize a large number of potential binding events such that they are spaced in 30bp intervals along the genome (Figure S13). Binding event positions are re-estimated over numerous EM training iterations, so that binding event discovery is not constrained by the initial 30bp interval (Figure S9). Alternatively, binding events can be initialized using the predicted peak positions of other peak callers, where potential binding events are initialized in 30bp intervals in a 500bp window around predicted peak positions. For example, ChExMix initial binding event positions in the MCF-7 analyses are initialized using MultiGPS results. The overall likelihood of the observed set of tags, *r*, given the binding event positions, *μ*, the binding event mixture probabilities (i.e. binding event strengths), *π*, and binding subtypes *τ* is defined as:

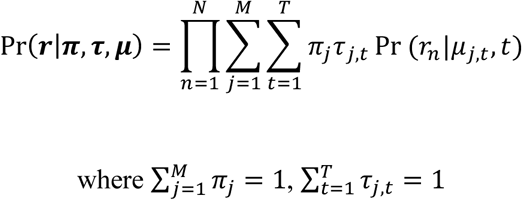

We incorporate biologically relevant assumptions in the form of priors on binding event strengths, binding locations, and subtype assignment. Similar to the GEM (Guo et al., 2012) and MultiGPS (Mahony et al., 2014) implementations, we place a sparseness promoting negative Dirichlet prior, *α*, on the binding strength *π* based on the assumption that binding events are relatively sparse throughout the genome (Neal and Hinton, 1998). We make two prior assumptions about binding subtype assignment: 1) the presence of subtype-specific DNA motif instances is indicative of the subtype to which a binding event belongs (i.e. can affect subtype probabilities); and 2) a binding event should be associated with a single subtype (i.e.
sparseness in subtype probabilities). To incorporate these assumptions, we place a Dirichlet prior *β* on the binding subtype probabilities *τ*.

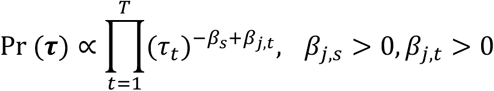

*β_s_* is the sparse prior parameter to adjust the degree of subtype sparseness:

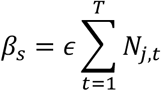

where *ϵ* is a parameter to tune the effect of the motif based prior, 0 ≤ *ϵ* ≤ 1. In this study, we choose *ϵ* = 0.05 (Figure S14, S15). *β_s_* is proportional to the number of tags assigned to the binding events.

*β_j,t_* denotes the binding subtype specific prior parameter and its value is proportional to *W_j,t_*, the strand specific log likelihood score for subtype *t*’s motif at event *j*’s location. *max W_j,t_* is the maximum possible log likelihood score from the weight matrix.

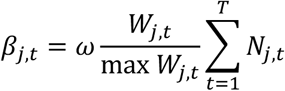

where *ω* is a parameter to tune the effect of the motif based prior, 0 ≤ *ω* ≤ 1. In this study, we choose *ω* = 0.2 (Figure S16). *N_j,t_* is the effective number of tags assigned to subtype *t* of the binding event *j*. The rationale is that a binding event *j* is more likely to be associated with subtype *t* if that subtype’s DNA motif is present in the vicinity. The parameter *β_j,t_* is scaled such that *β_j,t_* can be greater than *β_s_*. Therefore, a particular binding subtype will not be eliminated from consideration if the motif prior provides sufficient evidence of the binding subtype.

A positional prior on the base pair locations of binding events, *k*, is defined directly by subtype-specific motif log likelihood scores. Similar to MultiGPS, we introduce a Bernoulli prior over each genomic location where each element *k_i,t_* of the parameter *k* corresponds to the probability that genomic location *i* is a binding site of a binding type *t*. This prior assumes that there can be only one or zero binding events at a single position and that binding positions are selected independently along the genome according to this weighting. The positional prior is strand-specific. The prior assigns a likelihood to a set of binding sites on a genome of size *L* as follows:

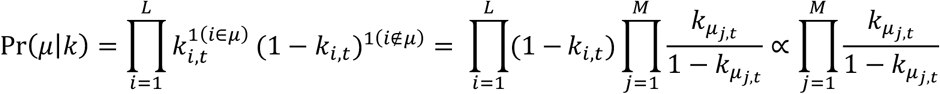

### Binding event prediction and subtype assignment

As in the original framework, the latent assignments of tags to binding events is represented by the vector *z*, where Pr(*z_n_* = *j*) = *π_j_*. The latent assignments of binding events to subtypes is represented by the vector *y*, where Pr(*y_j_* = *t*) = *τ_j,t_* The joint probability of latent variables is Pr(*z_n_* = *j, y_j_* = *t*) = *π_j_τ_j,t_*.

The complete-data log posterior is as follows:

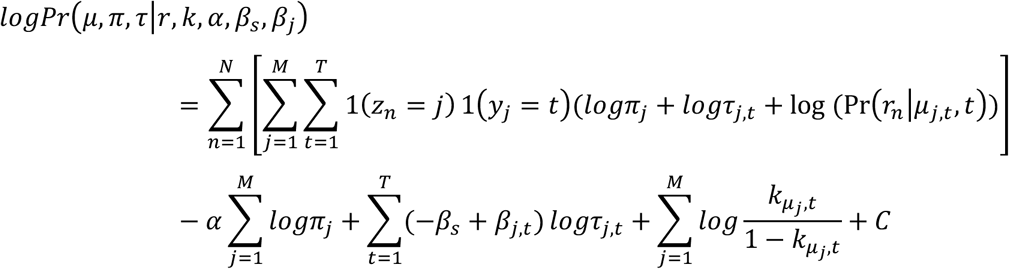

The overall binding event sparsity-inducing negative Dirichlet prior *α* acts only on the mixing probabilities *π*. Dirichlet priors *β_s_* and *β_j,t_* act only on the subtype probabilities *τ*. The positional prior acts only on the subtype binding event locations *μ*. The E-step thus calculates the relative responsibility of each binding subtype at each binding event in generating each tag as follows:

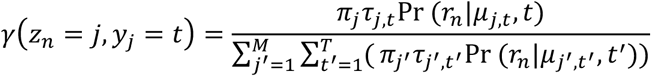

The maximum a posteriori probability (MAP) estimation (Figueiredo and Jain, 2002) of *π* and *τ* is as follows:

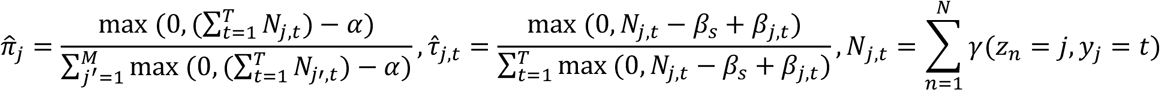

As in the MultiGPS framework, the *α* parameter can be interpreted as the minimum number of ChIP-exo tags required to support a binding event active in the model. Similarly, *β_s_* − *β_j,t_* is the minimum number of ChIP-exo tags required to support a binding event being associated with a particular binding subtype.

MAP values of *μ*_*j,t*_ are determined by enumerating over several possible values of *μ*_*j,t*_. Specifically, the MAP estimation of is:

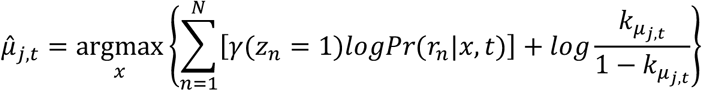

where *x* starts at the previous values of the position weighted by *τ* and expands outwards to 30bp each side. Each binding event is associated with a position weighted by subtype probabilities. If the maximization step results in two components sharing the same strand and weighted positions, they are combined in the next iteration of the algorithm.

As in our previous GPS frameworks, ChExMix requires that the number of tags associated with each predicted binding event be significantly higher than the scaled number of tags associated with the same binding events in a control experiment such as input or mock IP with exonuclease treatment (*p* < 0.001 with Benjamini-Hochberg corrected Binomial test). The control experiment normalization factors are estimated using the NCIS normalization method (Liang and Keles, 2012) with 10Kbp windows. Control tag counts are associated with individual binding events via maximum likelihood assignments using the trained model (i.e. assigning tags to binding events without changing model parameters such as *τ* or *μ*).

### Initial subtype characterization via tag distribution clustering

Subtypes may be initialized in ChExMix using tag distribution clustering, motif discovery, or a combination of both. To initialize subtypes via tag distribution clustering, we extract the stranded per-base tag counts in 150bp windows centered on the top 500 most enriched initial binding event positions. The per-base tag distributions are smoothed using a Gaussian kernel (variance = 1) and normalized by dividing by the sum of tag counts in the window. All pairs of binding event tag distributions are aligned against one another by finding the relative orientation and offset (in the range +/− 25bp) that produces the lowest Euclidean distance between normalized, smoothed tag distributions. Distances are converted to a pseudo-similarity score by multiplying by −1. Affinity propagation (Dueck and Frey, 2007) is applied to the similarity matrix (preference value = −0.1) to generate clusters. The number of clusters is automatically determined by the affinity propagation algorithm, albeit influenced by the preference value. Initial subtype-specific tag distributions are defined by the precomputed alignments against each cluster’s exemplar. During EM, subtype-specific tag distributions are updated by grouping binding events according to their maximum likelihood assigned subtypes and then combining each binding event’s assigned tag distributions.

### Initial subtype characterization via motif discovery

To characterize subtype-specific DNA motifs, ChExMix uses MEME (Bailey and Elkan, 1994) to discover a set of over-represented motifs in the top 1000 most enriched binding events (60bp windows). Motifs are retained if they discriminate bound regions from random sequences with true-positive vs. false-positive area under curve (AUC) above 0.7. Motif discovery is performed iteratively after removing the sequences containing previously discovered motifs until no further motifs pass the AUC threshold. Each discovered motif defines a subtype, and the corresponding tag distribution is defined using cumulative 5’ tag positions centered on motif instances within 30bp of binding events. Therefore, the number of motif-driven subtypes is determined by the number of motifs that pass the AUC threshold. When ChExMix is run with multiple ChIP-exo experiments, ChExMix performs a targeted motif discovery at sites where the predicted binding events from the two experiments occur within 30bp from each other. In this way, ChExMix attempts to identify unique motifs present in genomic regions where two proteins bind at proximal genomic loci.

### Merging initial subtypes and subtype re-estimation

If motif and tag distribution similarities from a pair of subtypes are above the thresholds (motif similarity using Pearson correlation > 0.95; tag distribution similarity using log KL divergence < −10), we retain only the subtype that is associated with the greater number of binding events. Subtypes are re-initialized during the second training iteration with the same approach. From the third training iteration, binding events are grouped into subtypes using maximum likelihood estimation and a targeted motif discovery is performed using the top 1000 most enriched subtype-specific binding events (60bp window). Subtypes are eliminated from the model during the subtype updates if the number of subtype-specific binding events falls below 5% of all binding events.

### Assessing subtype assignment performance using *in silico* mixed ChIP-exo data

To computationally simulate human ChIP-exo data that contains two distinct binding event subtypes, we mixed CTCF ChIP-exo data from HeLa cells (Rhee and Pugh, 2011), FoxA1 ChIP-exo data from MDA-MB-453 cells (Serandour et al., 2013), and an input control experiment from MCF-7 cells, all mapped to hg19. We first defined the top 20,000 binding event locations using MultiGPS for both CTCF and FoxA1 ChIP-exo experiments. We extended the binding events to 1Kbp regions and created a set of non-overlapping regions that contain peaks from either the CTCF or FoxA 1 experiment (but not both). To reflect the typical signal-to-noise ratio observed in real ChIP-exo experiments, 80% of the tags (24 million tags) come from the input control data, and the remaining (6 million) tags are randomly selected from all CTCF and FoxA1 ChIP-exo 1Kbp peak regions. We generated different datasets where the relative proportions of tags drawn from CTCF and FoxA1 experiments are varied. In these datasets, CTCF and FoxA1 ChIP-exo tags are always drawn randomly from all peak regions and are not preferentially drawn from the top-most binding events.

We ran the following binding event analysis methods on the simulation data: a) ChExMix with an option: --seqrmthres 0.3; b) ChExMix using default parameters with the exception of turning off the use of the motif prior in assigning subtypes (subtypes are still defined using motif discovery and tag distributions); and c) subtype assignment based on the ChExMix discovered motif hits. ChExMix recursively finds motifs by removing sequences with the previously discovered motifs. ChExMix option “--seqrmthres 0.3” (default value: 0.1) decreases the threshold to call motif hits to attempt to further deplete sequences with the previously discovered motifs. To scan ChExMix discovered motifs, we used the ChExMix discovered motifs to scan 60bp regions around all binding events and assigned subtypes based on the motif hits (log-likelihood scoring threshold of 3% per base FDR defined using a 2^nd^-order Markov model based on human genome nucleotide frequencies). Performance of binding subtype assignment is evaluated using labels based on whether the regions were taken from CTCF or FoxA1 ChIP-exo data. Sensitivity (TP/(TP+FN)) and specificity (TN/(TN+FP)) are used as the performance measures. The results show the average performance over five simulated datasets. We obtained CTCF and FoxA1 cognate DNA-binding motif rates (dashed lines in Figure 2) by scanning cis-bp database motifs (CTCF: M1957_1.02; FoxA1: M1965_1.02) (Weirauch et al., 2014) in 60bp regions around ChExMix peaks in the 100% CTCF and FoxA1 datasets, respectively, using 3% per base FDR.

### Performance of subtype discovery and classification in synthetic ChIP-exo data

To investigate ChExMix’s ability to learn and assign binding subtypes using only tag distribution information in a controlled setting, we used the ChIPReadSimulator module in SeqCode (https://github.com/seqcode/seqcode-core) to simulate two types of binding events using predefined ChIP-exo tag distributions. The tag distribution shapes used to define subtypes in these simulations (Figure 3A, 3B) were based on tag distributions observed in yeast Reb1 (subtype A) and human p53 (subtype B) ChIP-exo experiments (Reb1 and p53 distribution files available from https://github.com/seqcode/chexmix). We first simulated two datasets on a yeast-sized genome that consisted of pure signal; one of the datasets contained 500 subtype A binding events, while the other dataset contained 500 subtype B binding events. The relative strength of each of these binding events was drawn randomly from a distribution of relative tag counts observed for CTCF binding events in CTCF ChIP-seq experiments. Then, we modulated the relative sampling rate from each signal dataset and a background (mock IP control) dataset to create each individual simulated ChIP-exo dataset. Specifically, we varied the proportion of tags mixed between subtypes A and B to create different relative representations of binding event subtypes. We also modulated the proportions of tags drawn from the two signal experiments relative to that taken from the background (input) experiment. We ran ChExMix with the option “--nomotifs --scalewin 1000 --minmodelupdateevents 10”. Performance of binding subtype assignment is evaluated using 500bp window centered at simulated binding event locations. Sensitivity (TP/(TP+FN)) and specificity (TN/(TN+FP)) are used as the performance measures. Sensitivity and specificity reflect the accuracy of subtype assignments and are measured only with respect to detected binding events.

### Evaluating spatial resolution of ChIP-exo binding event predictions

To evaluate the spatial resolution performance of ChIP-exo peak callers, we quantify the distance between genomic coordinates of predicted binding events and high-scoring binding motif hits. As the center of the motif hit may not represent the true center of a binding event, we consider the distance between the predicted peaks to either edge of the motif. We compare spatial resolution on the set of predictions that are called by all methods and which have the same high-scoring motif hits (log-likelihood scoring threshold of 5% per base FDR defined using a 2^nd^-order Markov model based on the genomic nucleotide frequencies). Only events that occur within 50bp of a motif instance are included in the calculation. GEM is run with ChIP-exo specific parameters “--smooth 3 --mrc 20” as described in the documentation. MultiGPS is run with parameters “--fixedbp 20” with ChIP-exo tag distribution as described in the documentation. dFilter is run with a parameter “-ks 10” to decrease the kernel filter width. Q-nexus is run with parameters “-nexus-mode -s 100 -v” as described in the documentation. CexoR is run with parameters “idr=0.01, N=1e6, p=1e-9, dpeaks=c(0,150), dpairs=200” as suggested by the CexoR developer. All other software was run using default parameters.

### Public datasets

CTCF ChIP-exo in HeLa cells is obtained from SRA (SRA044886) and aligned against hg19 using Bowtie (Langmead et al., 2009) version 1.0.1 with options “-q --best --strata -m 1 --chunkmbs 1024 -C”. FoxA2 ChIP-exo in mouse liver is obtained from GEO (GSM1384738) and aligned against mm10 using BWA (Li and Durbin, 2009) version 0.6.2. FoxA1 ChIP-exo in MDA-MB-453 and input DNA in MCF-7 are downloaded from ERA (E-MTAB-1827) and aligned against hg19 using BWA version 0.7.12.

### ChIP-exo experiments and processing

The human breast adenocarcinoma cell line, MCF7, was obtained from American Type Culture Collection (ATCC) and cultured using DMEM with 10% heat inactivated FBS at 37°C with 5% CO_2_ in air. MCF7 cells were incubated in phenol red-free, charcoal stripped FBS for 48 hours prior to the 1 hour treatment with 17 *β*-estradiol (E2, Sigma) at 100 *μ*M. ChIP-exo assays for FoxA1, ERα, and CTCF were performed as previously described (Rhee and Pugh, 2011; Serandour et al., 2013). For ChIP-exo library preparation, affinity purified anti-FoxA1 (ab23738, Abcam; sc-514695 X, Santa Cruz), anti-ERα (ab108398, Abcam; sc8002 X, Santa Cruz), and anti-CTCF (07-729, Millipore) were incubated with chromatin. Mock IP control ChIP-exo experiments in MCF-7 cells were performed using the same approach but in the absence of antibody.

The *Saccharomyces cerevisiae* strain, BY4741, was obtained from Open Biosystems. Cells were grown in yeast peptone dextrose (YPD) media at 25°C to an OD_600_=0.8-1.0. Mock IP control ChIP-exo experiments in yeast were performed using rabbit IgG (Sigma, i5006) in the BY4741 background strain (which does not contain a tandem affinity purification tag sequence).

Libraries were paired-end sequenced and read pairs were mapped to the hg19 reference or sacCer3 genome using BWA version 0.7.12 with options “mem -T 30 -h 5”. Read pairs that share identical mapping coordinates on both ends are likely to represent PCR duplicates, and so Picard (http://broadinstitute.github.io/picard) was used to de-duplicate such pairs. Reads with MAPQ score less than 5 are filtered out using samtools (Li et al., 2009). During analysis of the MCF7 experiments, ChExMix was run with the following command-line parameters: --noclustering --q 0.05. ChExMix was initialized using the results of MultiGPS analysis of the dataset collection, where MultiGPS (version 0.74) was run using the following parameters: --q 0.05 --jointinmodel --fixedmodelrange --gaussmodelsmoothing --gausssmoothparam 1 --minmodelupdateevents 50.

## Availability

Open source code (MIT license) for ChExMix is available from https://github.com/seqcode/chexmix. Executables and documentation are available from http://mahonylab.org/software/chexmix. All ChIP-exo sequencing data produced in this study has been uploaded to GEO under accession GSE110502.

## Conflict of interest statement

BFP has a financial interest in Peconic, LLC, which utilizes the ChIP-exo technology implemented in this study and could potentially benefit from the outcomes of this research.

## Acknowledgements

The authors thank the members of the Center for Eukaryotic Gene Regulation at Penn State for helpful feedback and discussions.

## Funding

This manuscript is based upon work supported by the National Science Foundation ABI Innovation Grant No. DBI1564466 (to S.M.) Any opinions, findings and conclusions or recommendations expressed in this material are those of the authors and do not necessarily reflect the views of the National Science Foundation. This work was also supported by National Institutes of Health grant GM059055 (to B.F.P) and a Penn State Huck Graduate Research Innovation Award (to N.Y.).

## Supplement to Characterizing protein-DNA binding event subtypes in ChIP-exo data

### Robustness of ChExMix on various synthetic ChIP-exo data

We evaluated ChExMix performance on simulated ChIP-exo data that contain subtypes with either distinct motifs or tag distributions but not both. To simulate synthetic ChIP-exo data that have different motifs and the same tag distributions (Supplementary Figures S5 & S6), we selected 10,000 CTCF binding event locations for subtype A from CTCF ChIP-exo data in HeLa cells (Rhee and Pugh, 2011) and 10,000 FoxA1 binding event locations for subtype B from FoxA1 ChIP-exo data in MDA-MB-453 cells (Serandour et al., 2013). These selected locations thus contain motif instances associated with CTCF (subtype A) and FoxA1 (subtype B), respectively. In order to force the tag distributions at these binding locations to be equal, we simulated ChIP-exo binding events by randomly drawing ChIP-exo tag locations according to the yeast Reb1 ChIP-exo tag distribution (Figure 3A). The relative strength of each of these binding events (and thus the numbers of simulated tags at each binding event) was drawn randomly from a distribution of relative tag counts observed for CTCF binding events in CTCF ChIP-seq experiments. We then modulated the relative sampling rate from each signal dataset and a background (mock IP control) dataset to create each individual simulated ChIP-exo dataset with 3 million tags from either of the signal experiments and 27 million tags from the background. We varied the proportion of tags mixed between subtypes A and B to create different relative representations of binding event subtypes. We also used the same procedure to create a second set of simulated experiments containing two subtypes with different motifs and the same tag distribution, except this time we drew tags from the yeast Abf1 tag distribution (Reb1 and Abf1 distribution files available from https://github.com/seqcode/chexmix).

To generate synthetic ChIP-exo data that have same motifs and different tag distributions (Supplementary Figure S7), we selected the top 20,000 CTCF binding event locations in CTCF ChIP-exo data from HeLa cells (Rhee and Pugh, 2011). The CTCF binding event locations were randomly assigned to receive subtype A or B simulated binding events, such that one simulated dataset contained 10,000 subtype A binding events, while the other dataset contained 10,000 subtype B binding events. A high proportion of CTCF binding events contain the cognate binding motif, so therefore a large majority of the locations assigned to both subtypes will contain the same CTCF motif. At the selected locations, we simulated ChIP-exo binding events by randomly drawing ChIP-exo tag locations according to the yeast Reb1 (subtype A) or human p53 (subtype B) ChIP-exo tag distributions (Figure 3A, 3B). The relative strength of each of these binding events (and thus the numbers of simulated tags at each binding event) was drawn randomly from a distribution of relative tag counts observed for CTCF binding events in CTCF ChIP-seq experiments. We then modulated the relative sampling rate from each signal dataset and a background (mock IP control) dataset to create each individual simulated ChIP-exo dataset with 6 million tags from either of the signal experiments and 24 million tags from the background. We varied the proportion of tags mixed between subtypes A and B to create different relative representations of binding event subtypes. Performance of binding subtype assignment is evaluated using 500bp windows centered at simulated binding event locations. Sensitivity (TP/(TP+FN)) and specificity (TN/(TN+FP)) are used as the performance measures.

### Robustness of ChExMix on various read depths and replicates

To evaluate ChExMix performance on various read depths, we selected the CTCF and FoxA1 mixed dataset that has equal representations of subtypes (3 million CTCF tags and 3 million FoxA1 tags in 30 million total tags). We subsampled this ChIP-exo data to assess the sensitivity and specificity of subtype assignment by varying the read depth.

To evaluate the reproducibility of ChExMix results across true replicates, we used the replicate samples available in public data. For CTCF ChIP-exo, we obtained replicate 2 and replicate 3 from the accession number SRA044886. For FoxA1 ChIP-exo from MDA-MB-453 cells, we used a merge of replicate 2 & 4 and replicate 1 & 3 as two replicates from E-MTAB-1827, because individual replicates did not provide enough read coverage to generate the mixed ChIP-exo data. We followed the same procedures to create CTCF and FoxA1 mixed datasets with various subtype representations as described in the Methods section.

### ChExMix enables deconvolution of joint events

To examine ChExMix’s ability to resolve two closely spaced events, we simulated datasets by placing binding events at predefined intervals, and placed tags at those binding events by sampling from the ChIP-exo tag distribution observed in yeast Reb1 ChIP-exo experiments. We simulated a total of 40,000 binding events in a human-sized genome, but constrained the locations of 1000 events to occur within the range of 1 to 200bp from the neighboring events. The relative strength of each of these binding events was drawn randomly from a distribution of relative tag counts observed for CTCF binding events in CTCF ChIP-seq experiments. The simulation dataset contains 30 million tags. To reflect the typical signal-to-noise ratio observed in real ChIP-exo experiments, 80% of the tags are taken from the mock IP control. The remaining tags (6 million) are distributed among the binding events. We run ChExMix using default parameters with the exception of turning off the use of sequence information and the motif prior.

### Robustness of ChExMix to various initialization conditions

We examine the performance of ChExMix on different initialization conditions. During the initialization of binding events, ChExMix places binding components every 30 base pairs. We analyze how different spacing of components affects the sensitivity of peak detection and running time of the algorithm. We computationally mixed tags from CTCF ChIP-exo and input background, using an approach similar to the *in silico* mixed CTCF FoxA1 ChIP-exo experiment described in the Methods section. We created simulation data by drawing 6 million CTCF tags from 1Kbp regions centered around CTCF binding events from CTCF ChIP-exo data and 24 million background tags from the input control. ChExMix is run with an option “--noflanking”. This option will ensure that ChExMix will not automatically place additional binding components during the EM iterations. ChExMix performance is evaluated based on sensitivity of recovering predefined peak locations. We score peaks as positive if ChExMix peaks occur within 50bp of MultiGPS peak locations. The results show that ChExMix stably recovers above 90% of peaks when component spacing intervals are smaller than 100 base pair (Figure S13). The sensitivity drops significantly when the component intervals become bigger than 200 base pairs.

### Sparsity and motif prior weights in subtype assignment

In this section, we examined the effect of varying the sparsity and motif prior weights on subtype assignment. We assume that binding events should be associated with a single subtype. Hence, we employ a sparseness promoting prior in assigning binding subtypes to encourage a single subtype to dominate the probabilities. In assessing the effect of varying this prior, we used simulated ChIP-exo data that mix equal proportions of CTCF and FoxA1 ChIP-exo tags (as described in Methods). We used the F1 score to measure the performance of subtype assignment, calculated as the following using the scikit-learn python package:

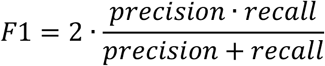

The results show that ChExMix performance drops significantly when the sparsity prior is above 0.1 (Figure S14). We observe equal representations of each subtype when we increase the sparseness promoting prior above 0.1. Subtype probability distributions shift towards 1 as we change the sparseness promoting prior to 0, 0.05, and 0.1 (Figure S15). The current default of 0.05 shifts the maximum assignment probability distribution towards 1 with a minor decrease in performance. Hence, we use 0.05 as the default value of the subtype sparsity prior.

Next, we evaluated how different motif weights affect ChExMix performance using the same CTCF/FoxA1 mixed data. Motif weights control the balance between tag distribution and sequence in subtype assignment. We measure the performance using F1 score as described above. We observe that performance continues to increase as the motif prior increases (Figure S16). Our current motif prior default is 0.2 because we do not wish sequence information to dominate subtype assignment.

**Figure S1.**
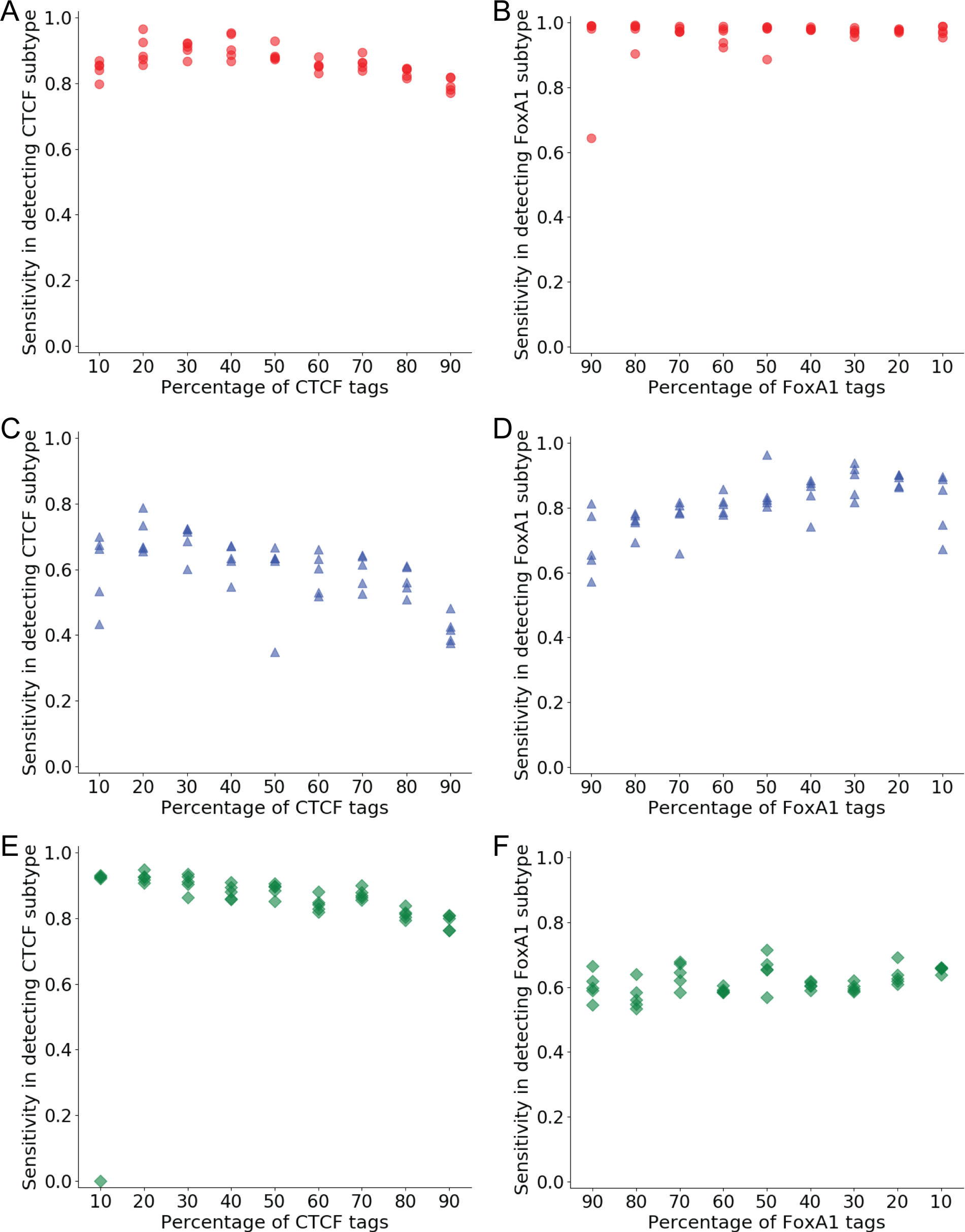
Related to Figure 2C, D. Sensitivity in subtype assignment from five simulation datasets. Plots show sensitivity for correctly assigning binding events to the CTCF and FoxA1 subtypes using ChExMix with *de novo* estimated tag distributions and motifs (A, B), ChExMix with tag distributions alone (C, D), and scanning ChExMix discovered motifs (E, F). The relative proportion of signal tags is varied between the CTCF and FoxA1 experiments.

**Figure S2.**
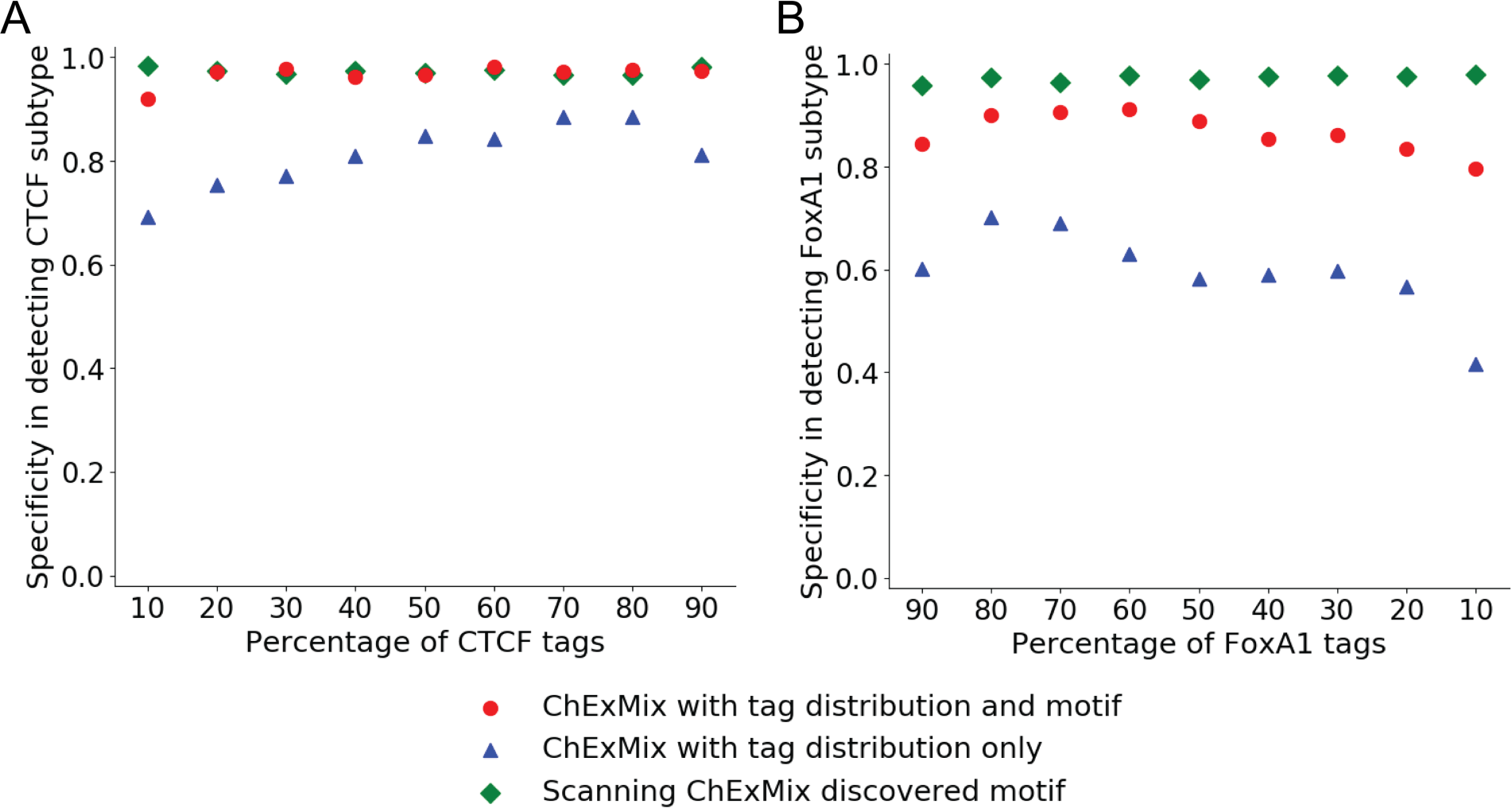
Related to Figure 2C, D. Specificity in subtype assignment using ChExMix with *de novo* estimated tag distributions and motifs (red dots), ChExMix with tag distributions alone (blue triangles), and scanning ChExMix discovered motifs (green diamonds). Plots show specificity for correctly assigning binding events to the CTCF (A) and FoxA1 (B) subtypes, which varies as the relative proportion of signal tags is varied between the CTCF and FoxA1 experiments. Each data point represents an average performance of five simulations.

**Figure S3.**
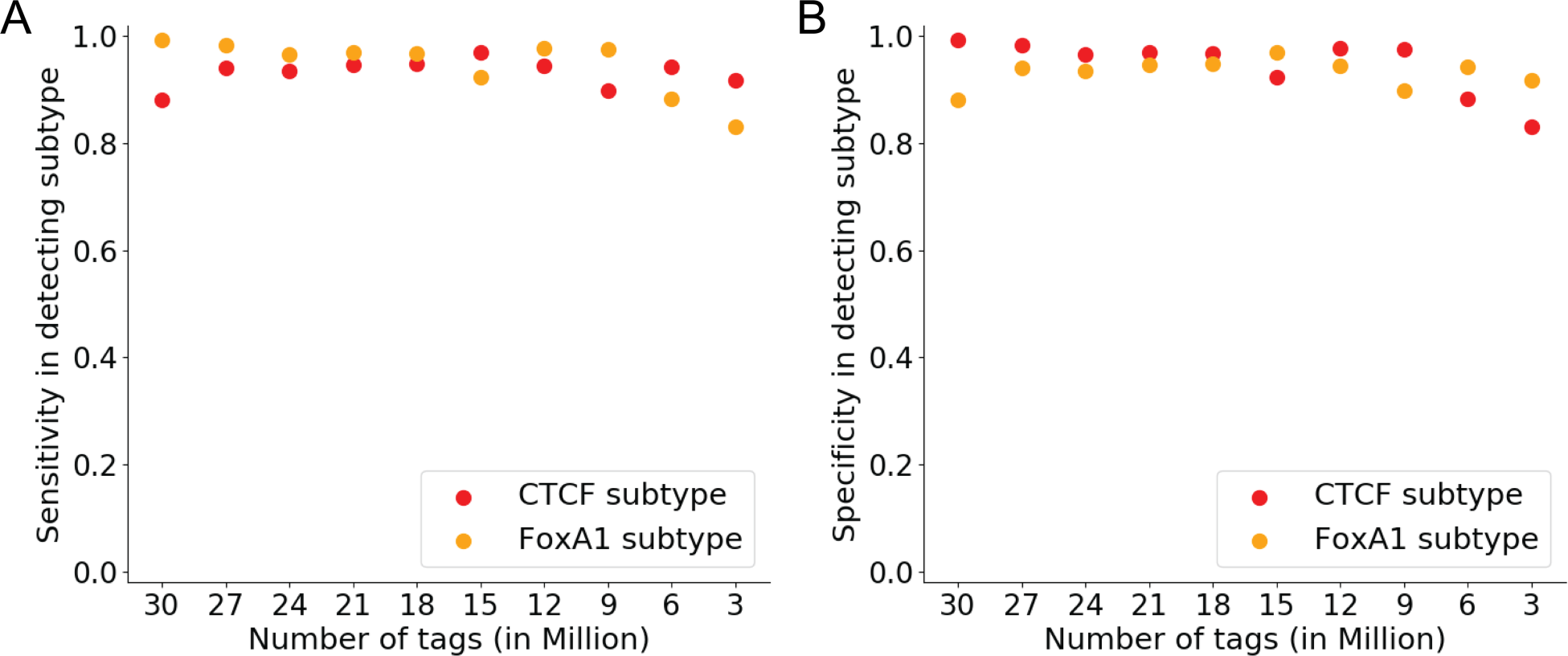
Sensitivity (A) and specificity (B) of subtype assignment across various read depths of simulation datasets. Plots show sensitivity for correctly assigning binding events to the CTCF subtype (red dots) and FoxA1 subtype (orange dots), as the read depth of the experiments is varied.

**Figure S4.**
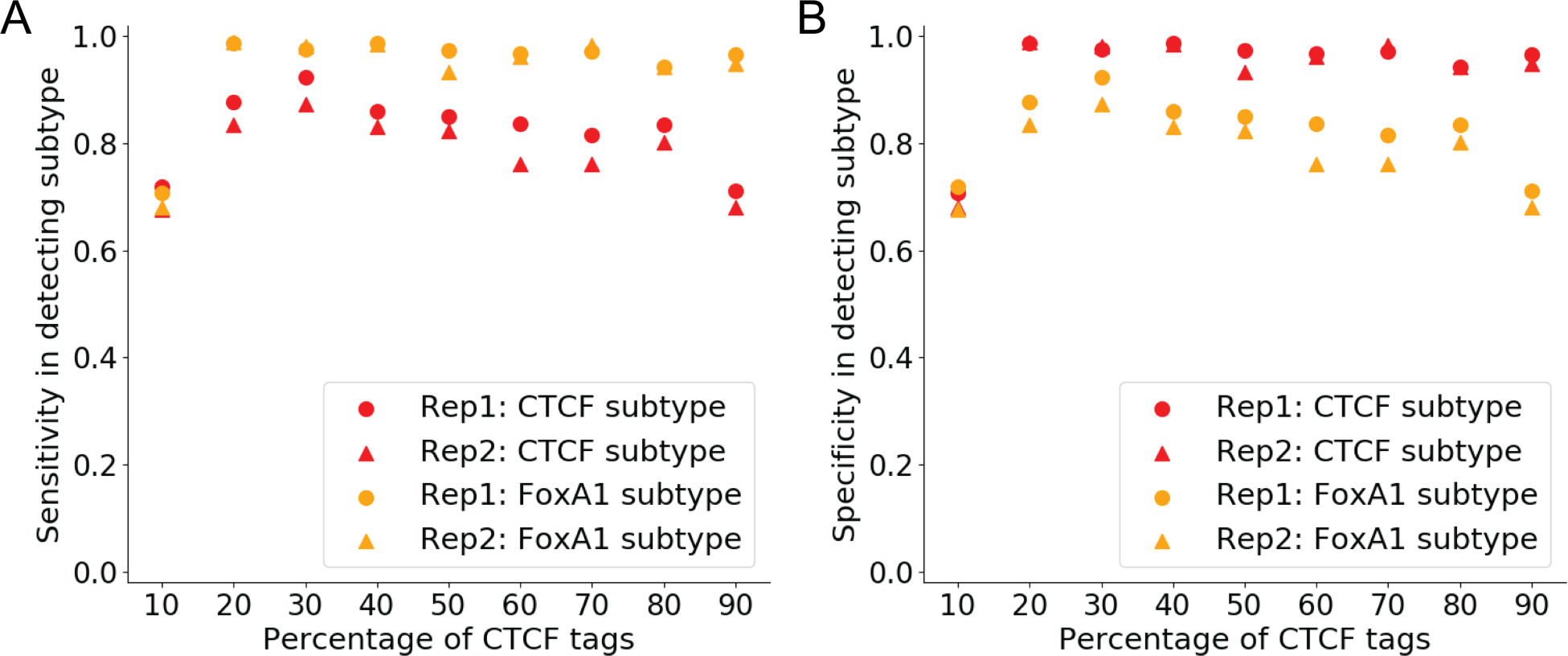
Sensitivity (A) and specificity (B) of subtype assignment from biological replicates of simulation datasets. Plots show sensitivity/specificity for correctly assigning binding events to the CTCF subtype in replicate 1 (red dots) & replicate 2 (red triangles) and FoxA1 subtype in replicate 1 (orange dots) & replicate 2 (orange triangles), as the relative proportion of signal tags is varied between the two subtypes.

**Figure S5.**
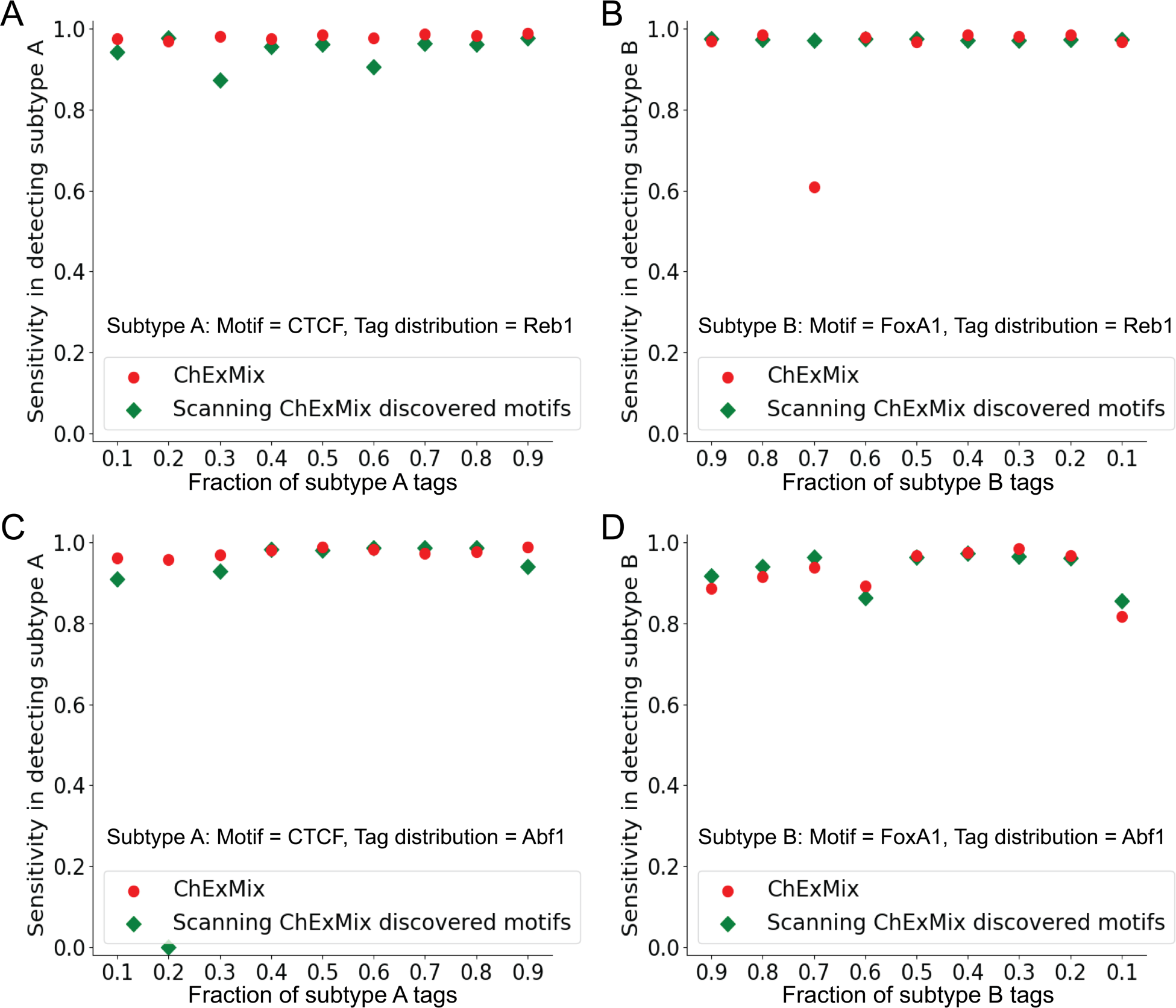
Sensitivity in subtype assignment with simulated datasets that have different motifs and the same tag distribution. Plots show sensitivity for correctly assigning binding events to subtype A (CTCF motif) and subtype B (FoxA1 motif) (red dots) with Reb1 distribution (A, B) and Abf1 distribution (C, D) and *de novo* estimated motif instances alone (green diamonds) (A-D), as the relative proportion of signal tags is varied between the two subtypes.

**Figure S6.**
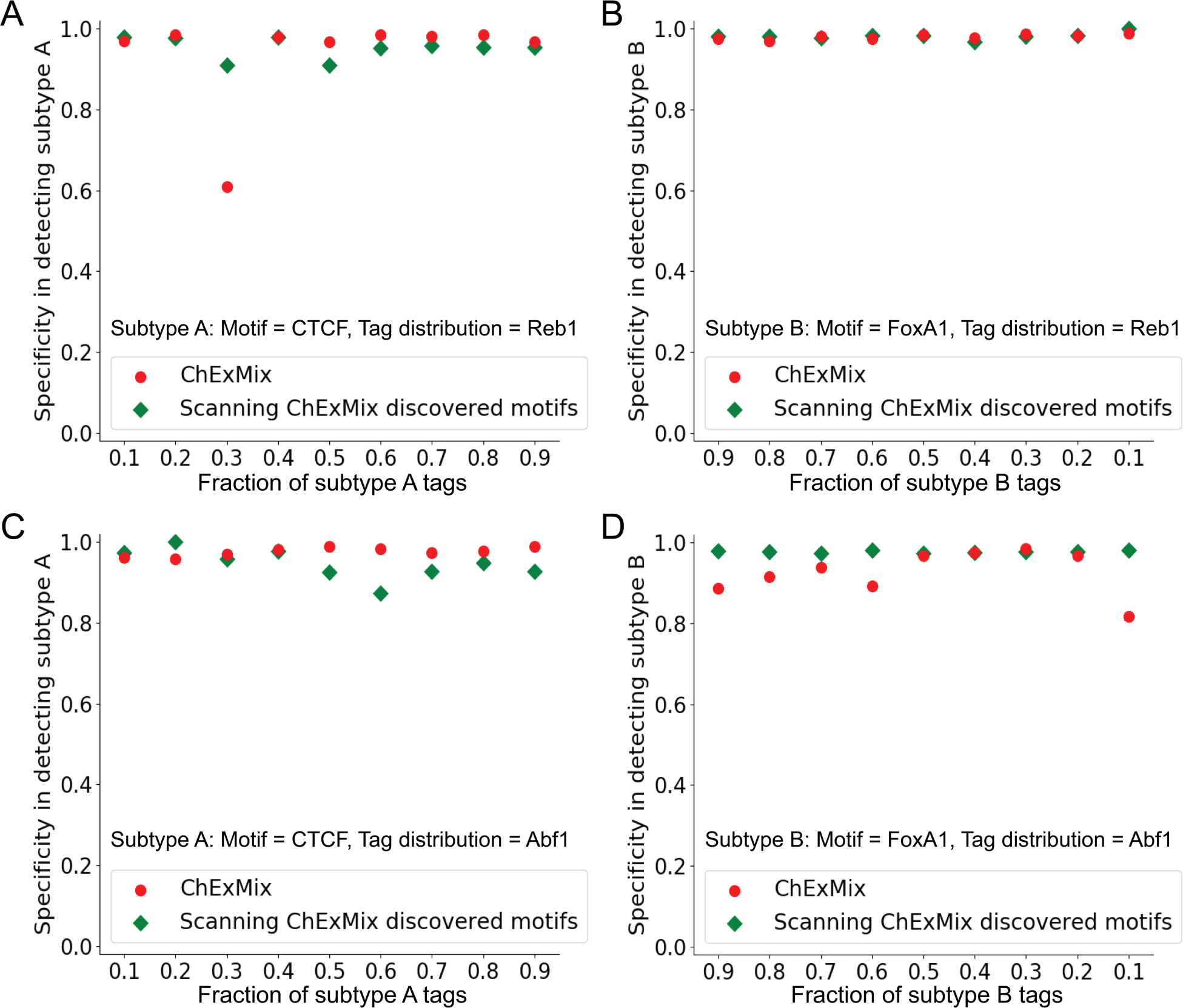
Specificity in subtype assignment with simulated datasets that have different motifs and the same tag distribution. Plots show specificity for correctly assigning binding events to subtype A (CTCF motif) and subtype B (FoxA1 motif) (red dots) with Reb1 distribution (A, B) and Abf1 distribution (C, D) and *de novo* estimated motif instances alone (green diamonds) (A-D), as the relative proportion of signal tags is varied between the two subtypes.

**Figure S7.**
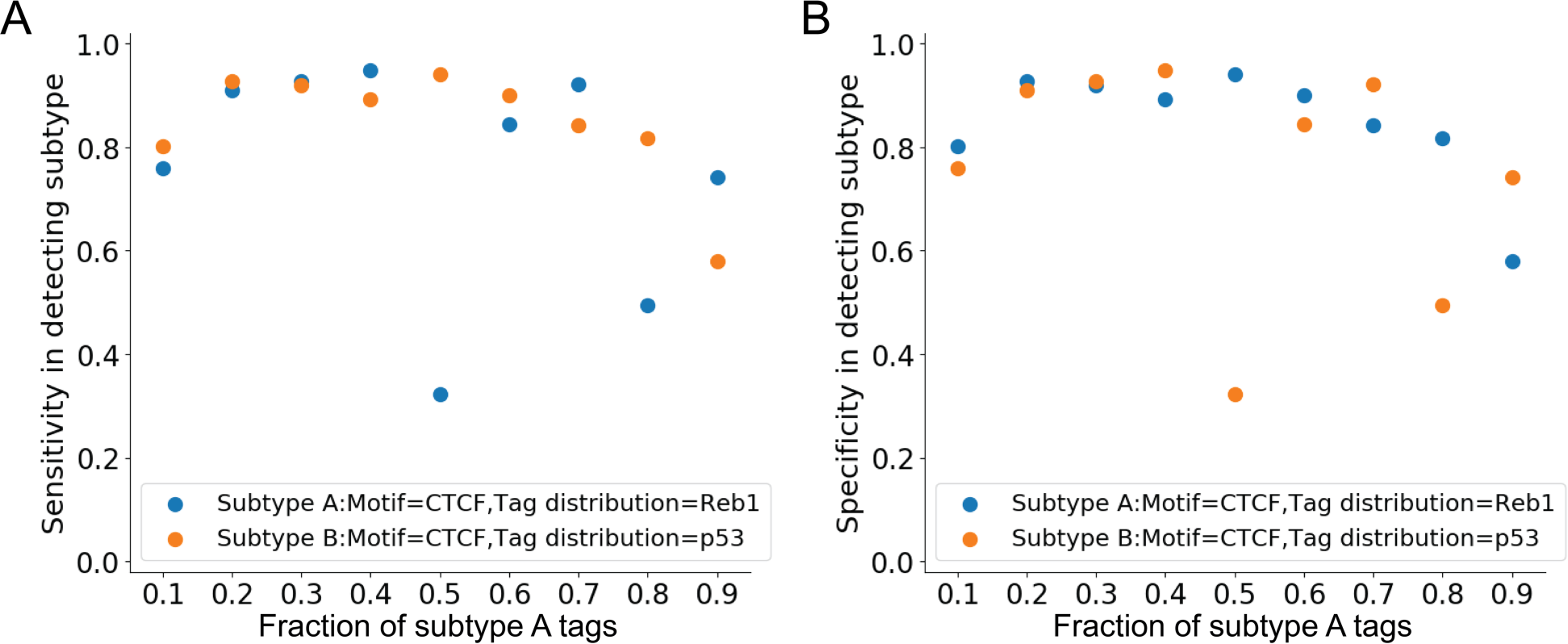
Sensitivity (A) and specificity (B) in subtype assignment from simulation data with the same motif and different tag distributions. Plots show sensitivity/specificity for correctly assigning binding events to the subtype A (Reb1 distribution) (blue dots) and subtype B (p53 distribution) (orange dots), as the relative proportion of signal tags is varied between the two subtypes.

**Figure S8.**
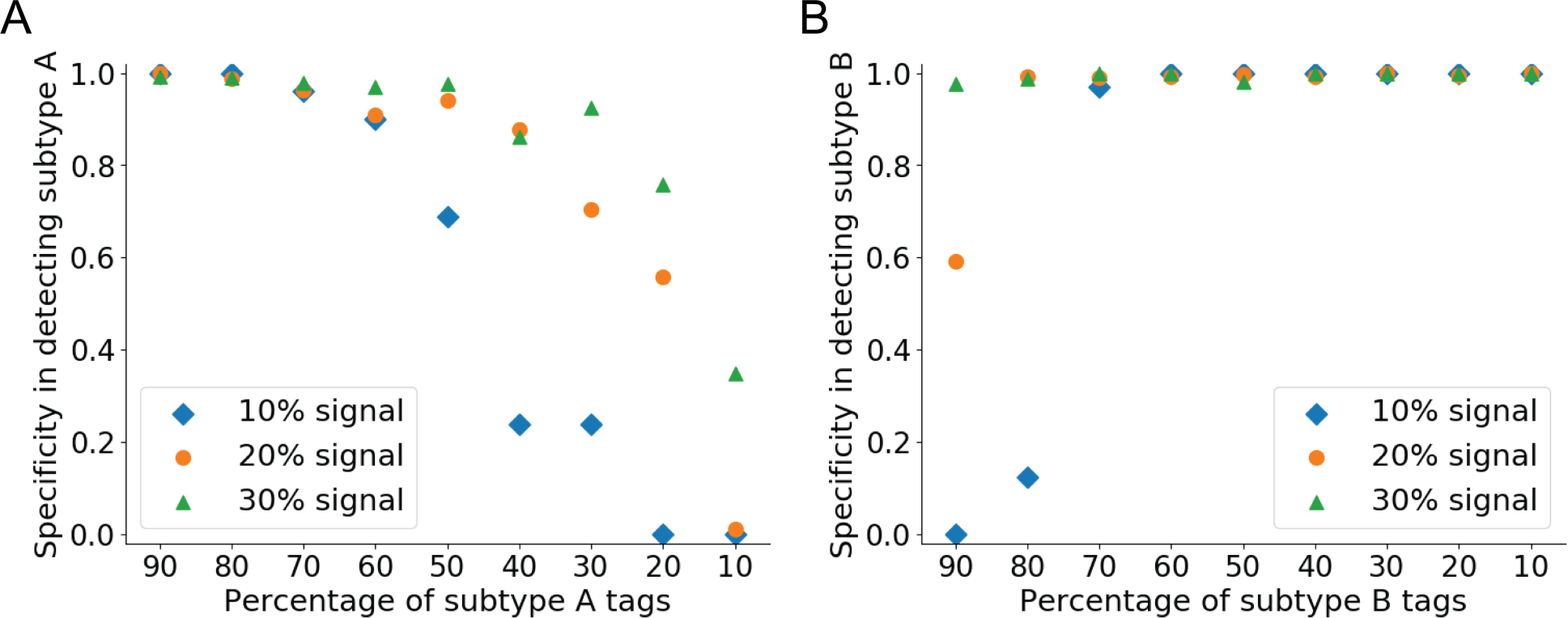
Related to Figure 3G, H. Specificity in subtype assignment using *de novo* estimated tag distributions with overall signal of 10% (blue diamonds), 20% (orange dots), and 30% (green triangles). Plots show specificity for correctly assigning binding events to the subtype A (Reb1 distribution) (A) and subtype B (p53 distribution) (B) subtypes, as the relative proportion of signal tags is varied between the two subtypes.

**Figure S9.**
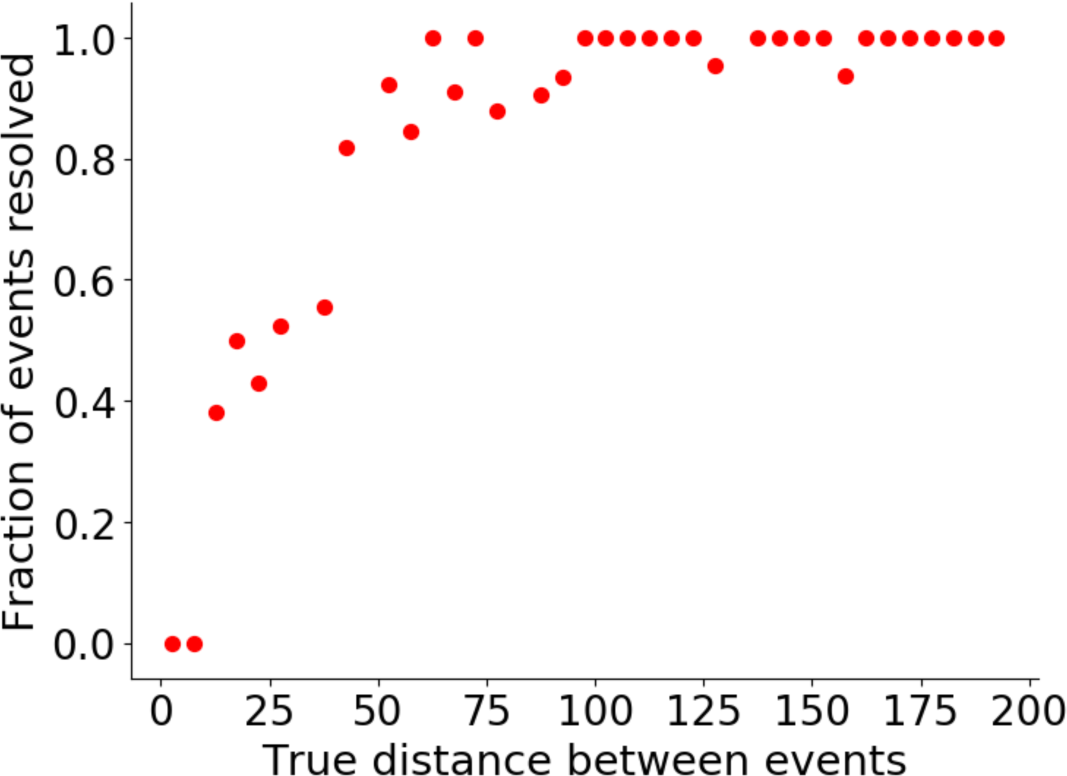
ChExMix is able to resolve closely spaced binding events. Joint events are placed between a range of 1 to 200bp apart from each other. The x-axis shows the true distance between events. The y-axis shows the fraction of the events resolved to be two binding events. The results are averaged every 5bp.

**Figure S10.**
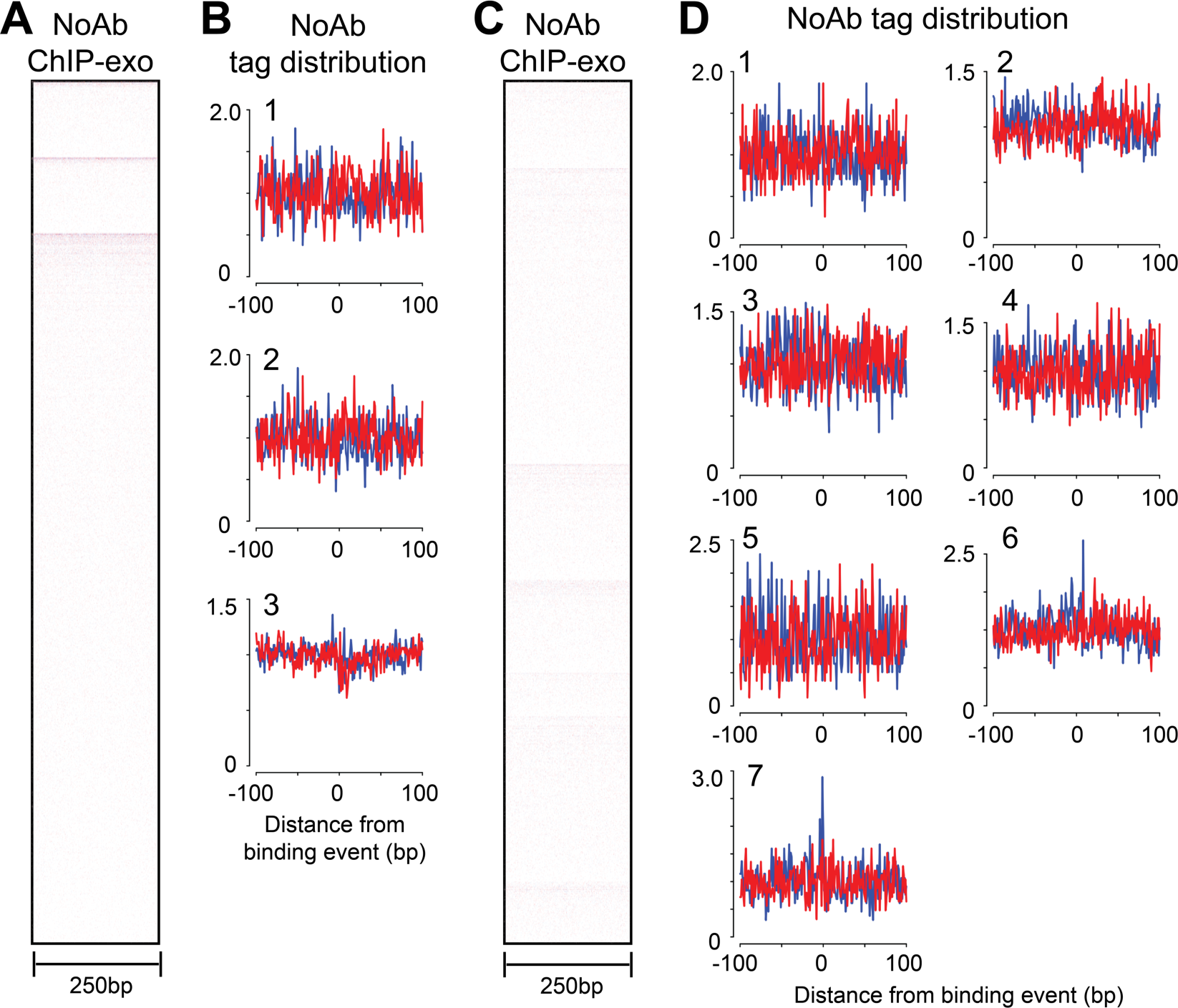
Heatmap and tag distributions of no antibody control ChIP-exo at FoxA1 and ERα ChIP-exo binding events. A) Heat map of no antibody control ChIP-exo tags and B) tag distributions at FoxA1 subtype 1, 2 and 3 binding events. C) Heat map of no antibody control and D) tag distributions at ERα subtypes 1 to 7.

**Figure S11.**
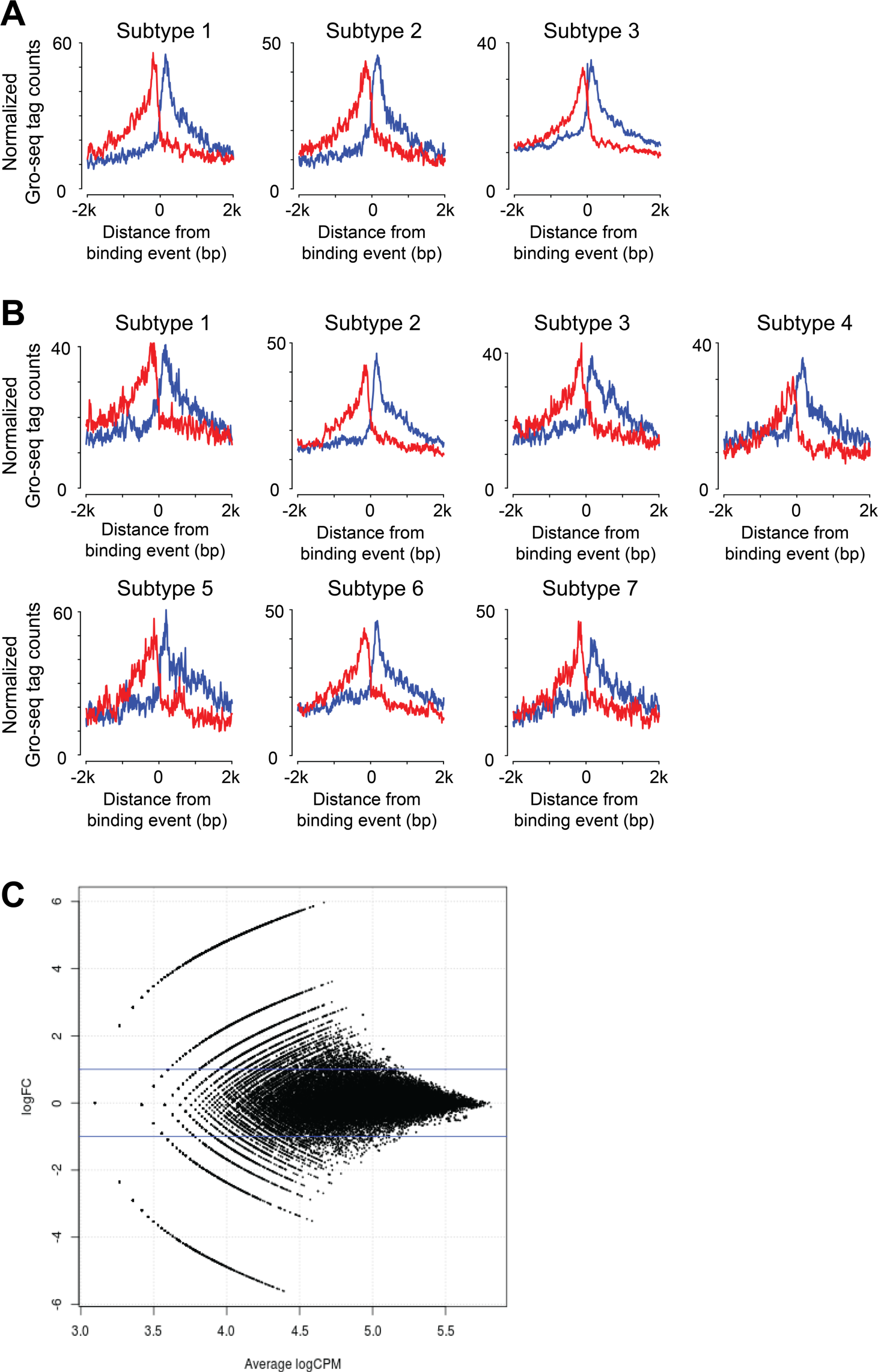
GRO-seq analysis at ERα and FoxA1 ChIP-exo subtypes. A) Normalized GRO-seq tag counts at three FoxA1 subtypes in estradiol treated MCF-7 cells. B) Normalized GRO-seq tag counts at seven ERα subtypes in estradiol treated MCF-7 cells. C) MA plot showing GRO-seq tag counts in a 4Kbp window at ERα and FoxA1 binding events, where the y-axis represents the tag count ratio between estradiol treated and untreated, and the x-axis represents the mean tag count. No ERα and FoxAl binding events displayed significant differential enrichment for GRO-seq tags (p<0.001).

**Figure S12.**
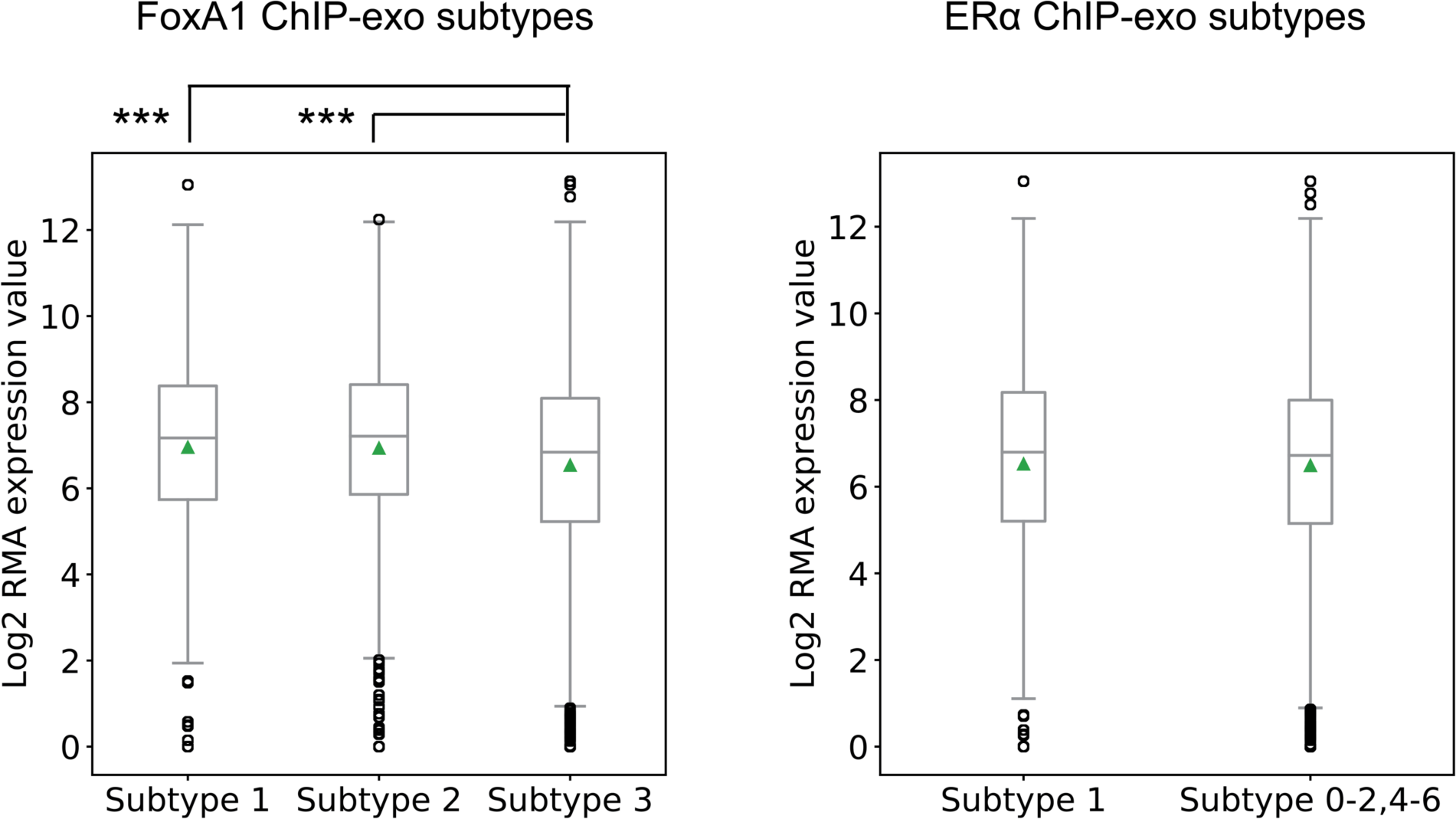
Microarray analysis of FoxA1 and ERα ChIP-exo subtypes. We used GREAT to associate binding events with up to two nearest genes that occur within 10Kbp (McLean et al., 2010). Microarray gene expression of subtype 1 (with ERα like motif) and subtype 2 (with CTCF like motif) are significantly higher than subtype 3 in FoxA1 ChIP-exo (*** p-value < 0.001). No significant differences are detected between subtype 4 (with a Forkhead motif) and the rest of the ERα ChIP-exo subtypes.

**Figure S13.**
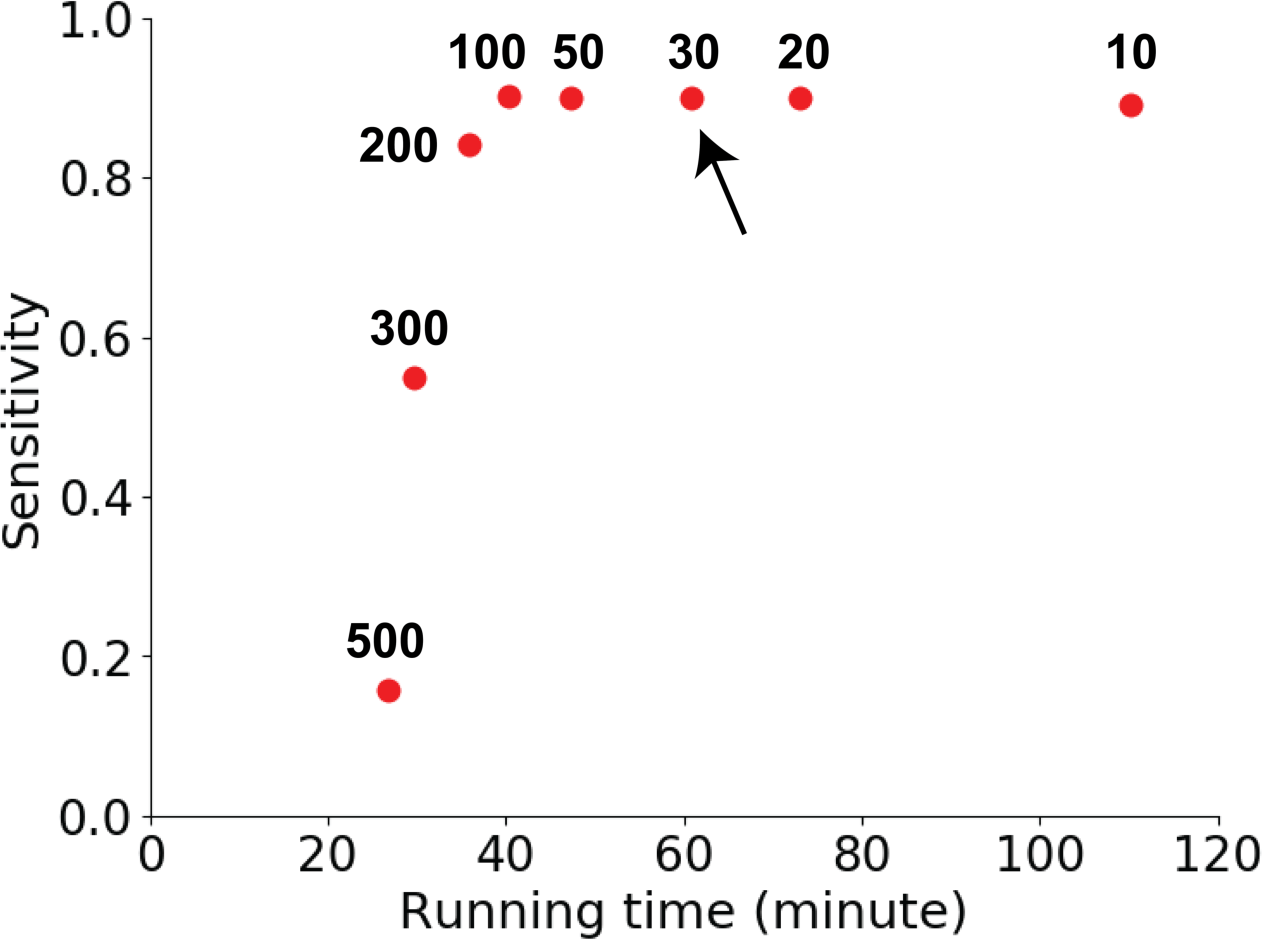
ChExMix sensitivity in detecting ChIP-exo peaks with various initialization conditions. Potential binding event mixture components are placed in intervals; 10, 20,30, 50, 100, 200, 300, and 500bp. Performance of ChExMix is evaluated by the percentage of true peaks recovered and the running time of the algorithm. The current default value is 30 bp (black arrow).

**Figure S14.**
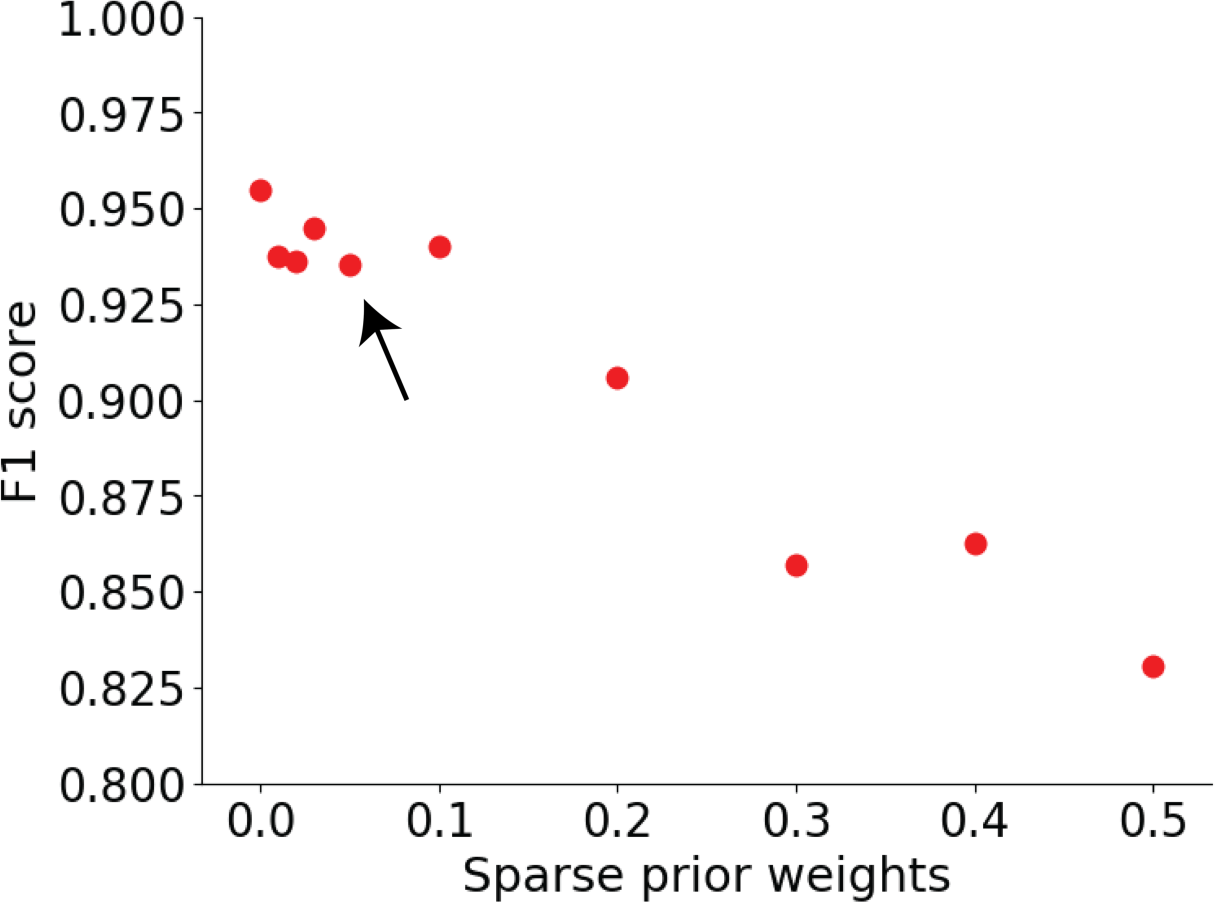
Effect of varying sparseness promoting prior weight in ChExMix performance. Sparse prior weights are varied between 0 to 0.5. F1 score is used to evaluate the performance of subtype assignment. The current default of sparse prior weight is 0.05 (black arrow).

**Figure S15.**
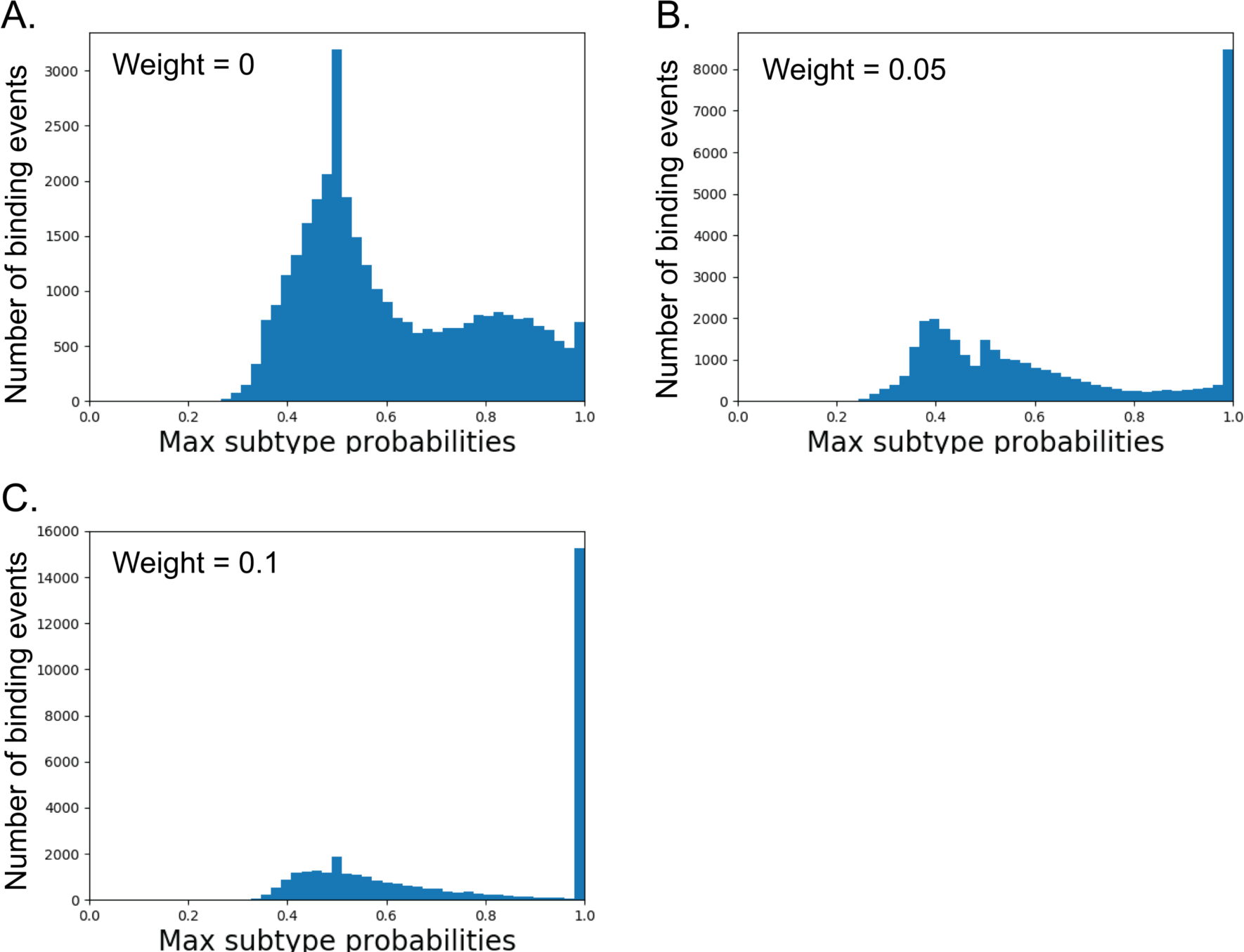
Effect of varying sparseness promoting prior weight in maximum subtype probabilities. Plots show the distributions of maximum subtype probabilities at the motif weight of 0 (A), 0.05 (B; the current default), and 0.1 (C).

**Figure S16.**
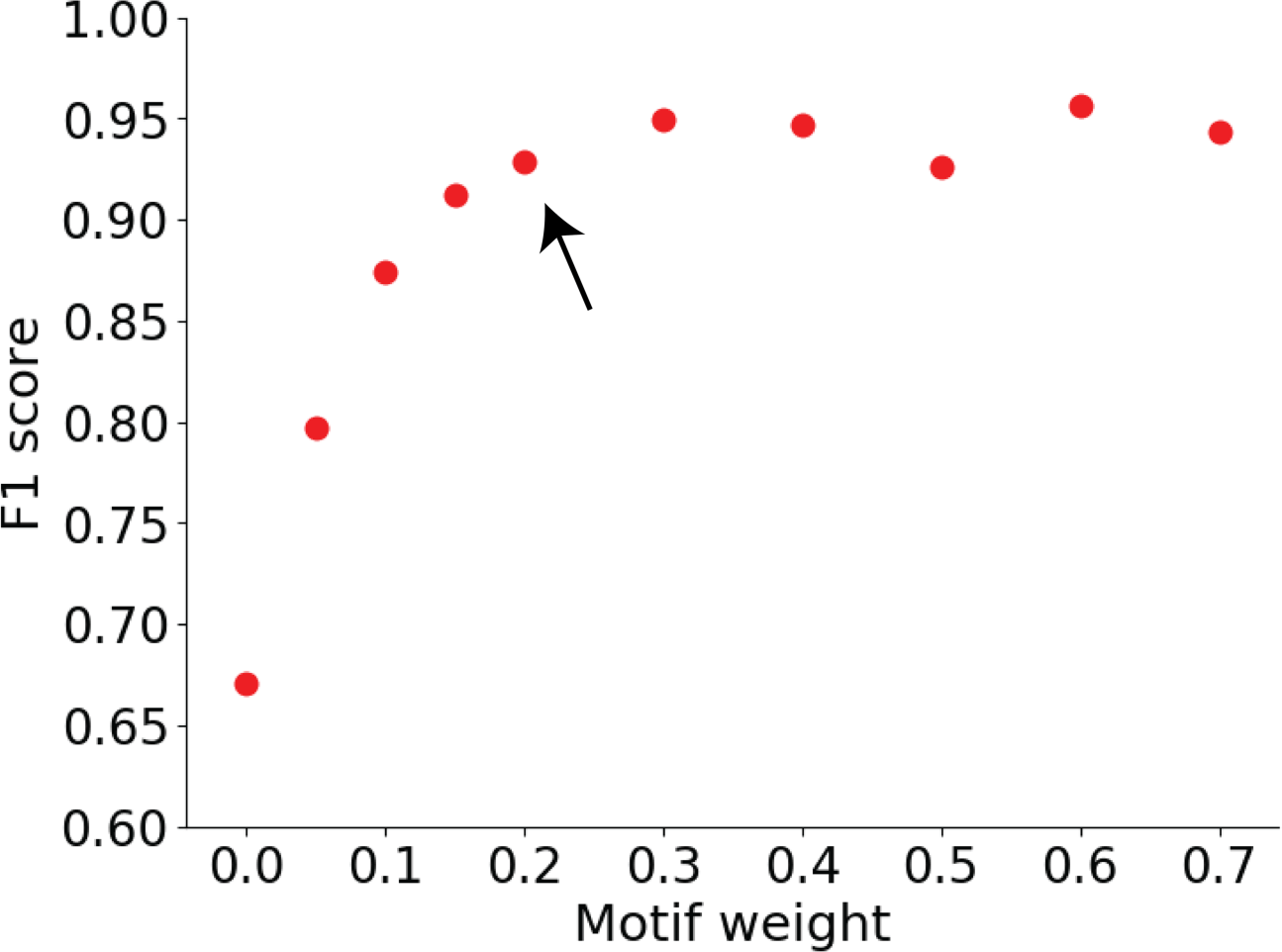
Effect of varying motif prior weight in ChExMix performance. The motif weights are varied between 0 to 0.7. F1 score is used to evaluate the performance in subtype assignment. The current default value is 0.2 (black arrow).

**Table S1.**
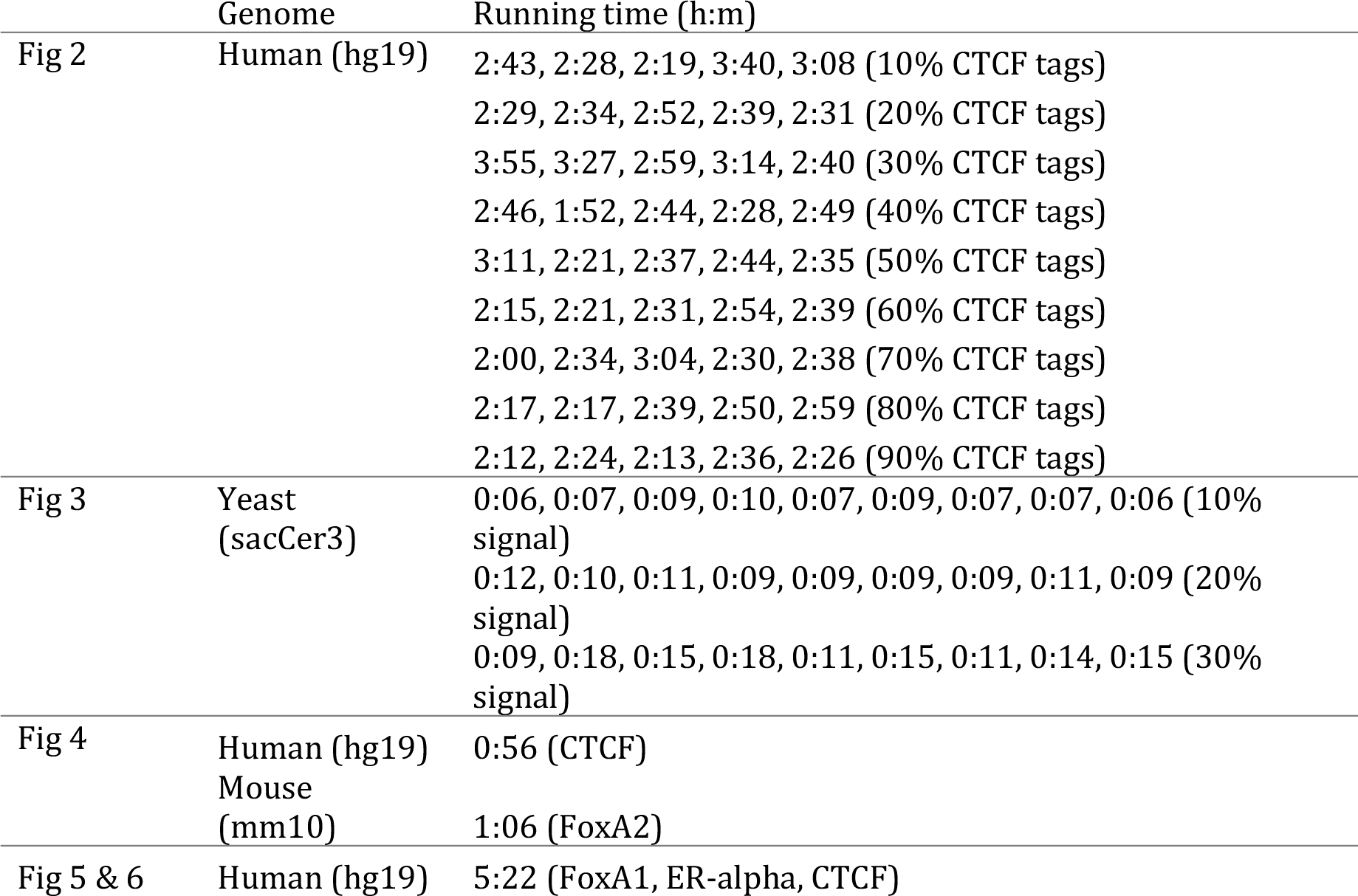
ChExMix running time. Jobs were run on an Intel Xeon E5-2680 v2 2.8GHz server blade using 20 threads.

|  | Genome | Running time (h:m) |
| --- | --- | --- |
| Fig 2 | Human (hg19) | 2:43, 2:28, 2:19, 3:40, 3:08 (10% CTCF tags)<br>2:29, 2:34, 2:52, 2:39, 2:31 (20% CTCF tags)<br>3:55, 3:27, 2:59, 3:14, 2:40 (30% CTCF tags)<br>2:46, 1:52, 2:44, 2:28, 2:49 (40% CTCF tags)<br>3:11, 2:21, 2:37, 2:44, 2:35 (50% CTCF tags)<br>2:15, 2:21, 2:31, 2:54, 2:39 (60% CTCF tags)<br>2:00, 2:34, 3:04, 2:30, 2:38 (70% CTCF tags)<br>2:17, 2:17, 2:39, 2:50, 2:59 (80% CTCF tags)<br>2:12, 2:24, 2:13, 2:36, 2:26 (90% CTCF tags) |
| Fig 3 | Yeast (sacCer3) | 0:06, 0:07, 0:09, 0:10, 0:07, 0:09, 0:07, 0:07, 0:06 (10% signal)<br>0:12, 0:10, 0:11, 0:09, 0:09, 0:09, 0:09, 0:11, 0:09 (20% signal)<br>0:09, 0:18, 0:15, 0:18, 0:11, 0:15, 0:11, 0:14, 0:15 (30% signal) |
| Fig 4 | Human (hg19)<br>Mouse (mm10) | 0:56 (CTCF)<br>1:06 (FoxA2) |
| Fig 5 & 6 | Human (hg19) | 5:22 (FoxA1, ER-alpha, CTCF) |

**Table S2.** Related to Figure 2C, D. Number of subtypes discovered from five simulation datasets. The table shows the number of CTCF- and FoxA1-related subtypes discovered by ChExMix in each simulated dataset when using *de novo* estimated tag distributions and motifs (1st & 2nd columns) or using tag distributions alone (3rd & 4th columns). The relative proportion of signal tags is varied between the CTCF and FoxA1 experiments.

| % CTCF tags | ChExMix with de novo estimated tag distributions and motifs |  | ChExMix with tag distributions alone |  |
| --- | --- | --- | --- | --- |
|  | CTCF subtype | FoxA1 subtype | CTCF subtype | FoxA1 subtype |
| 10 | 1, 1, 1, 2, 1 | 4, 3, 1, 4, 5 | 1, 1, 1, 2, 1 | 4, 3, 1, 4, 5 |
| 20 | 1, 3, 2, 3, 2 | 4, 1, 1, 3, 1 | 1, 2, 1, 2, 2 | 4, 2, 2, 4, 1 |
| 30 | 3, 2, 2, 3, 3 | 3, 2, 4, 3, 3 | 3, 1, 2, 3, 3 | 3, 3, 4, 3, 3 |
| 40 | 3, 1, 3, 1, 3 | 1, 1, 1, 1, 1 | 2, 1, 2, 1, 2 | 2, 1, 2, 1, 2 |
| 50 | 3, 3, 3, 3, 3 | 3, 3, 3, 1, 3 | 3, 3, 3, 2, 3 | 3, 3, 3, 2, 3 |
| 60 | 2, 1, 2, 2, 2 | 3, 2, 3, 3, 4 | 2, 1, 1, 2, 2 | 3, 2, 4, 3, 4 |
| 70 | 2, 2, 3, 3, 4 | 2, 3, 4, 2, 2 | 1, 2, 3, 2, 3 | 3, 3, 4, 3, 3 |
| 80 | 2, 2, 5, 2, 2 | 2, 2, 3, 3, 3 | 2, 1, 5, 2, 2 | 2, 3, 3, 3, 3 |
| 90 | 2, 2, 1, 3, 2 | 2, 3, 2, 2, 2 | 1, 3, 1, 3, 2 | 3, 2, 2, 2, 2 |

**Table S3.**
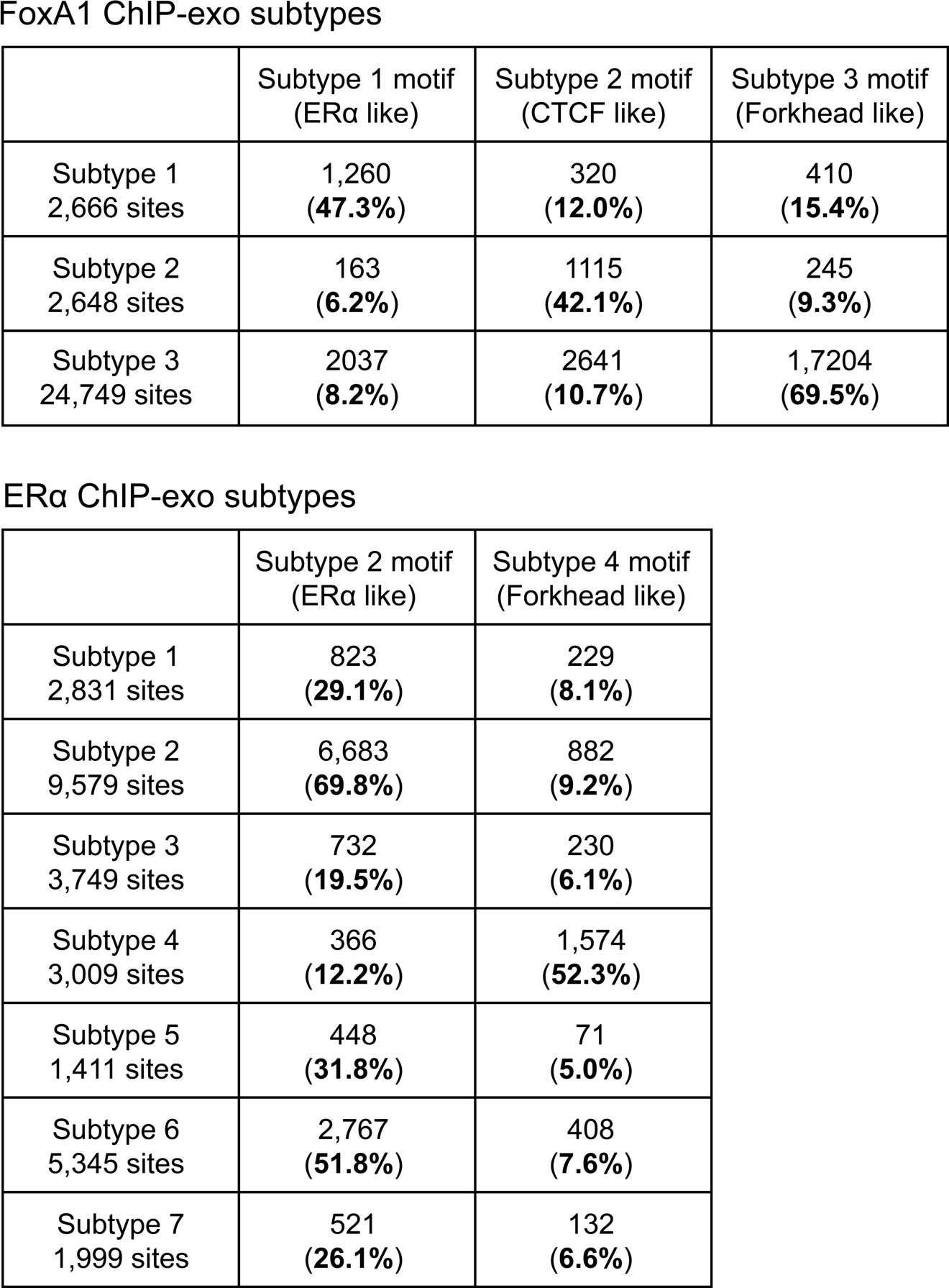
Subtype-specific motif occurrence rates in FoxA1 and ERα ChIP-exo binding events. Subtype-specific motifs are scanned in a 100bp window around subtype-specific binding events (5% per base FDR using a Markov model based on human nucleotide frequencies).

FoxA1 ChIP-exo subtypes
| | Subtype 1 motif<br>(ERα like) | Subtype 2 motif<br>(CTCF like) | Subtype 3 motif<br>(Forkhead like) |
| --- | --- | --- | --- |
| Subtype 1<br>2,666 sites | 1,260<br>(**47.3%**) | 320<br>(**12.0%**) | 410<br>(**15.4%**) |
| Subtype 2<br>2,648 sites | 163<br>(**6.2%**) | 1115<br>(**42.1%**) | 245<br>(**9.3%**) |
| Subtype 3<br>24,749 sites | 2037<br>(**8.2%**) | 2641<br>(**10.7%**) | 1,7204<br>(**69.5%**) |

**Table S3.** Subtype-specific motif occurrence rates in FoxA1 and ERα ChIP-exo binding events. Subtype-specific motifs are scanned in a 100bp window around subtype-specific binding events (5% per base FDR using a Markov model based on human nucleotide frequencies).
| | Subtype 2 motif<br>(ERα like) | Subtype 4 motif<br>(Forkhead like) |
| --- | --- | --- |
| Subtype 1<br>2,831 sites | 823<br>(**29.1%**) | 229<br>(**8.1%**) |
| Subtype 2<br>9,579 sites | 6,683<br>(**69.8%**) | 882<br>(**9.2%**) |
| Subtype 3<br>3,749 sites | 732<br>(**19.5%**) | 230<br>(**6.1%**) |
| Subtype 4<br>3,009 sites | 366<br>(**12.2%**) | 1,574<br>(**52.3%**) |
| Subtype 5<br>1,411 sites | 448<br>(**31.8%**) | 71<br>(**5.0%**) |
| Subtype 6<br>5,345 sites | 2,767<br>(**51.8%**) | 408<br>(**7.6%**) |
| Subtype 7<br>1,999 sites | 521<br>(**26.1%**) | 132<br>(**6.6%**) |

